# High-quality chromosome-scale assembly of the walnut (*Juglans regia* L) reference genome

**DOI:** 10.1101/809798

**Authors:** Annarita Marrano, Monica Britton, Paulo A. Zaini, Aleksey V. Zimin, Rachael E. Workman, Daniela Puiu, Luca Bianco, Erica Adele Di Pierro, Brian J. Allen, Sandeep Chakraborty, Michela Troggio, Charles A. Leslie, Winston Timp, Abhaya Dandekar, Steven L. Salzberg, David B. Neale

**Affiliations:** Department of Plant Sciences, University of California, Davis, CA 95616, USA; Bioinformatics Core Facility, Genome Center, University of California Davis, CA 95616, USA; Department of Biomedical Engineering, Johns Hopkins University, Baltimore, MD 21205, USA; Center for Computational Biology, Whiting School of Engineering, Johns Hopkins University, Baltimore, MD 21205, USA; Research and Innovation Center, Department of Genomics and Biology of Fruit Crops, Fondazione E Mach, San Michele all’ Adige (TN) 38010, Italy; Departments of Computer Science and Biostatistics, Johns Hopkins University, Baltimore, MD 21218

**Keywords:** Nanopore, Hi-C, IsoSeq, gene prediction, genetic diversity, proteome, allergens

## Abstract

The release of the first reference genome of walnut (*Juglans regia* L.) enabled many achievements in the characterization of walnut genetic and functional variation. However, it is highly fragmented, preventing the integration of genetic, transcriptomic, and proteomic information to fully elucidate walnut biological processes. Here we report the new chromosome-scale assembly of the walnut reference genome (Chandler v2.0) obtained by combining Oxford Nanopore long-read sequencing with chromosome conformation capture (Hi-C) technology. Relative to the previous reference genome, the new assembly features an 84.4-fold increase in N50 size, and the full sequence of all 16 chromosomal pseudomolecules, nine of which present telomere sequences at both ends. Using full-length transcripts from single-molecule real-time sequencing, we predicted 40,491 gene models, with a mean gene length higher than the previous gene annotations. Most of the new protein-coding genes (90%) are full-length, which represents a significant improvement compared to Chandler v1.0 (only 48%). We then tested the potential impact of the new chromosome-level genome on different areas of walnut research. By studying the proteome changes occurring during catkin development, we observed that the virtual proteome obtained from Chandler v2.0 presents fewer artifacts than the previous reference genome, enabling the identification of a new potential pollen allergen in walnut. Also, the new chromosome-scale genome facilitates in-depth studies of intraspecies genetic diversity by revealing previously undetected autozygous regions in Chandler, likely resulting from inbreeding, and 195 genomic regions highly differentiated between Western and Eastern walnut cultivars. Overall, Chandler v2.0 is a valuable resource to understand and explore walnut biology better.

## INTRODUCTION

Persian walnut (*Juglans regia* L.) is among the top three most-consumed nuts in the world, and over the last ten years, its global production increased by 37% (International Nut and Dried Fruit Council, 2019). Its richness in alpha-linolenic acid (ALA), proteins, minerals and vitamins along with documented benefits for human health explains this increased interest in walnut consumption (Martínez et al. 2010). As suggested by its generic name *Juglans* from the Latin appellation ‘*Jovis glans*’, which loosely means ‘nut of gods’, the culinary and medical value of Persian walnut was already widely prized by ancient civilizations (McGranahan and Leslie 2012).

The origin and evolution of the Persian walnut are the results of a complex interplay between hybridization, human migration and biogeographical forces (Pollegioni et al. 2017). A recent phylogenomic analysis revealed that Persian walnut (and its landrace *J. sigillata*) arose from an ancient hybridization between American black walnuts and Asian butternuts during the late Pliocene (3.45 Mya) (Zhang et al. 2019). Evidence suggests that the mountains of Central Asia were the cradle of domestication of Persian walnut (Zeven and Zhukovskiĭ 1975), from where it spread to the rest of Asia, the Balkans, Europe and, finally, the Americas.

Today, walnut is cultivated worldwide in an area of 1,587,566 ha, mostly in China and the USA (FAOSTAT statistics, 2017). Considerable phenotypic and genetic variability can be observed in this wide distribution area, especially in the Eastern countries, where walnuts can still be found in wild fruit forests. Many studies on genetic diversity in walnut have outlined a genetic differentiation between Eastern and Western genotypes (Ebrahimi et al. 2016; Marrano et al. 2018). Moreover, walnuts from Eastern Europe, Central Asia, and China exhibit higher genetic diversity and a higher number of rare alleles than the genotypes from Western countries (Bernard et al. 2018a).

The release of the first reference genome, Chandler v1.0 (Martínez-García et al. 2016), enabled the study of walnut genetics at a genome-wide scale. For the first time, it was possible to explore the gene space of Persian walnut with the prediction of 32,498 gene models, providing the basis to untangle complex phenotypic pathways, such as those responsible for the synthesis of phenolic compounds. The availability of a reference genome marked the beginning of a genomics phase in Persian walnut, allowing whole-genome resequencing (Stevens et al. 2018; Zhang et al. 2019), the development of high-density genotyping tools (Marrano et al. 2018; Kefayati et al. 2018) and the genetic dissection of important agronomical traits in walnut (Arab et al. 2019; Famula et al. 2019; Marrano et al. 2019). However, the Chandler v1.0 assembly is highly fragmented, compromising the accuracy of gene prediction and the fulfillment of advanced genomics studies necessary to resolve many, still unanswered questions in walnut research.

The recent introduction of long-read sequencing technologies and long-range scaffolding methods has enabled chromosome-scale assembly for multiple plant species, including highly heterozygous tree crops such as almond (*Prunus dulcis*; (Sánchez-Pérez et al. 2019) and kiwifruit (*Actinidia eriantha*; (Tang et al. 2019). The availability of genomes with fully assembled chromosomes provides foundations for understanding plant domestication and evolution (Jarvis et al. 2017; Maccaferri et al. 2019; Sánchez-Pérez et al. 2019), the mechanisms governing important traits (e.g. flower color and scent; (Raymond et al. 2018), as well as the impact of epigenetic modifications on phenotypic variability (Daccord et al. 2017). Recently, Zhu et al., (2019) assembled the parental genomes of a hybrid *J. microcarpa* × *J. regia* (cv. Serr) at the chromosome-scale using long-read PacBio sequencing and optical mapping. They relied on the haplotype divergence between the two *Juglans* species and demonstrated an ongoing asymmetric fractionation of the two subgenomes present in *Juglans* genomes.

Here we report a new chromosome-level assembly of the walnut reference genome with unprecedented contiguity, Chandler v2.0, which we obtained by combining Oxford Nanopore long-read sequencing (Lu et al. 2016) with chromosome conformation capture (Hi-C) technology (Belton et al. 2012). Thanks to the increased contiguity of Chandler v2.0, we were able to substantially improve gene prediction accuracy, with new, longer gene models identified and many fewer artifacts compared to Chandler v1.0. Also, the availability of full, chromosomal sequences reveals new genetic diversity of Chandler, previously inaccessible through standard genotyping tools, and significant genetic differentiation between Western and Eastern walnuts at 195 genomic regions, including also loci involved in nut shape and harvest date. In the present research, we demonstrate the fundamental role of a chromosome-scale reference genome to integrate transcriptomics, population genetics, and proteomics, which in turn enable a better understanding of walnut biology.

## RESULTS AND DISCUSSION

### Genome long-read sequencing and assembly

To increase the contiguity of the Chandler genome, we first generated deep sequence coverage using Oxford Nanopore Technology (ONT), a cost-effective long-read sequencing approach that determines DNA bases by measuring the changes in electrical conductivity generated while DNA fragments pass a tiny biological pore (Leggett and Clark 2017). Since the release of the first plant genome assembly generated using ONT sequencing (Schmidt et al. 2017), this technology has been applied to sequence and obtain chromosome-scale genomes of many other plant species (Belser et al. 2018; Yasodha et al. 2018; Deschamps et al. 2018). In Persian walnut, ONT sequencing yielded 7,096,311 reads that provided 21.9 Gbp of sequence, or ∼35X genome coverage (assuming a genome size of 620 Mb). Read lengths averaged 3.1 kb, and the N50 read length was 6.7 kb, with the longest read being 992.2 kb.

One of the major limitations of long-read sequencing technologies is their high error rate, which can range between 5% and 15% for Nanopore sequencing (Rang et al. 2018). To overcome this limitation, we adopted the hybrid assembly technique incorporated into the MaSuRCA assembler, which combines long, high-error reads with shorter but much more accurate Illumina sequencing reads to generate a robust, highly contiguous genome assembly (Zimin et al. 2017). First, using the Illumina reads, we created 3.7 million ‘super-reads’ with a total length of 2.9 Gb. We then combined the super-reads with the ONT reads to generate 3.2 million mega-reads with a mean length of 4.7 kb, representing 24X genome coverage (**Supplemental Table S1**). Finally, we assembled the mega-reads to obtain the ‘hybrid’ Illumina-ONT assembly, which comprised 1,498 scaffolds, 258 contigs, and 25,007 old scaffolds from Chandler v1.0 (**Table 1; Supplemental Table S2**).

**Table 1.**
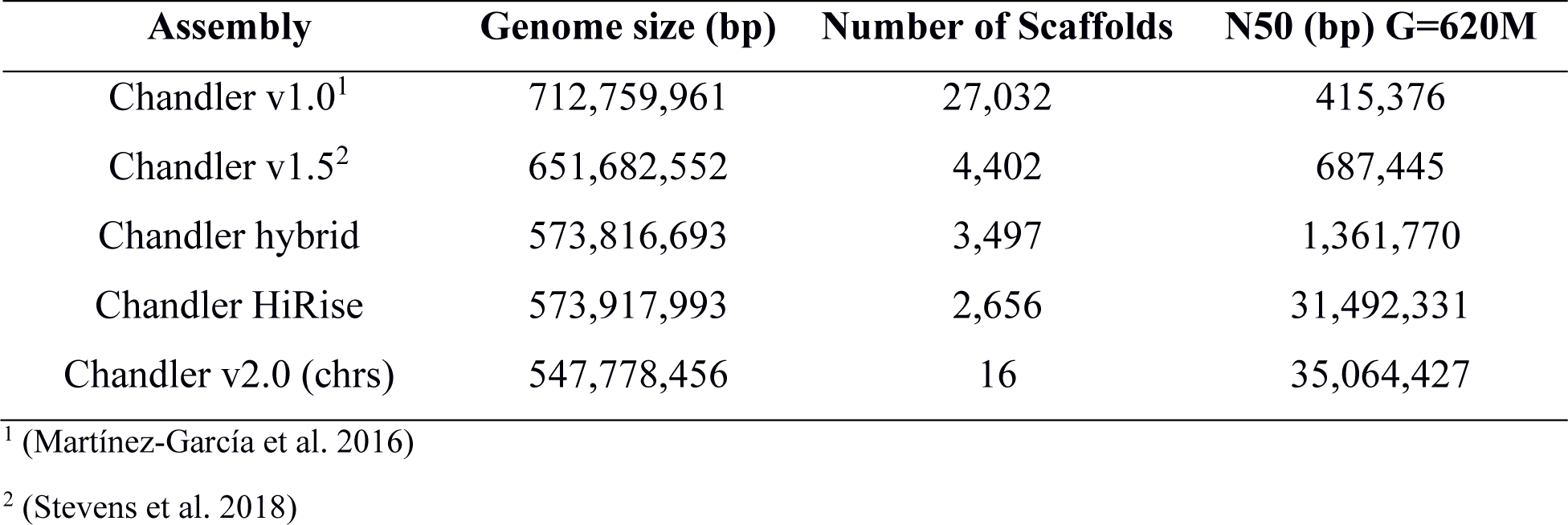
Comparison among the four assemblies of Chandler. Scaffolds shorter than 1,000 bp are not included in these totals.

| Assembly | Genome size (bp) | Number of Scaffolds | N50 (bp) G=620M |
| --- | --- | --- | --- |
| Chandler v1.0<sup>1</sup> | 712,759,961 | 27,032 | 415,376 |
| Chandler v1.5<sup>2</sup> | 651,682,552 | 4,402 | 687,445 |
| Chandler hybrid | 573,816,693 | 3,497 | 1,361,770 |
| Chandler HiRise | 573,917,993 | 2,656 | 31,492,331 |
| Chandler v2.0 (chrs) | 547,778,456 | 16 | 35,064,427 |

<sup>1</sup> (Martínez-García et al. 2016)
<sup>2</sup> (Stevens et al. 2018)

Even though the total number of scaffolds (> 1 Kb) was reduced by 80% compared to Chandler v1.0 (**Table 1**), the new hybrid assembly was still fragmented. To improve the assembly further and build chromosome-scale scaffolds, we applied Hi-C sequencing, which is based on proximity ligation of DNA fragments in their natural conformation within the nucleus (Belton et al. 2012). The HiRise scaffolding pipeline processed 356 million paired-end 100-bp Illumina reads to generate the HiRise assembly, which contained 2,656 scaffolds longer than 1 kb (**Table 1**). The top 17 scaffolds from this assembly spanned more than 90% of the total assembly length, with a scaffold length ranging from 19.6 to 45.2 Mb (**Supplemental Figure S1-S2**). As compared to the previous (1.0) assembly, the Chandler genome contiguity increased dramatically, with an N50 size 98% higher than Chandler v1.0 and only 0.04% of the genome in gaps.

### Validation of the HiRise assembly

To assess the quality of the HiRise assembly, we used two independent sources of data. First, we used the single nucleotide polymorphism (SNP) markers mapped on the high-density genetic map of Chandler recently described by (Marrano et al. 2019). Out of the 8,080 SNPs mapped into 16 linkage groups (LGs), 6,894 had probes aligning uniquely on the HiRise assembly with 98% of identity for more than 95% of their length. A total of 35 scaffolds of the HiRise assembly could be anchored to a chromosomal linkage group by at least one SNP (**Figure 1**). In particular, 13 LGs were spanned by a single HiRise scaffold, while two to three scaffolds each aligned the remaining three LGs.

**Figure 1.**
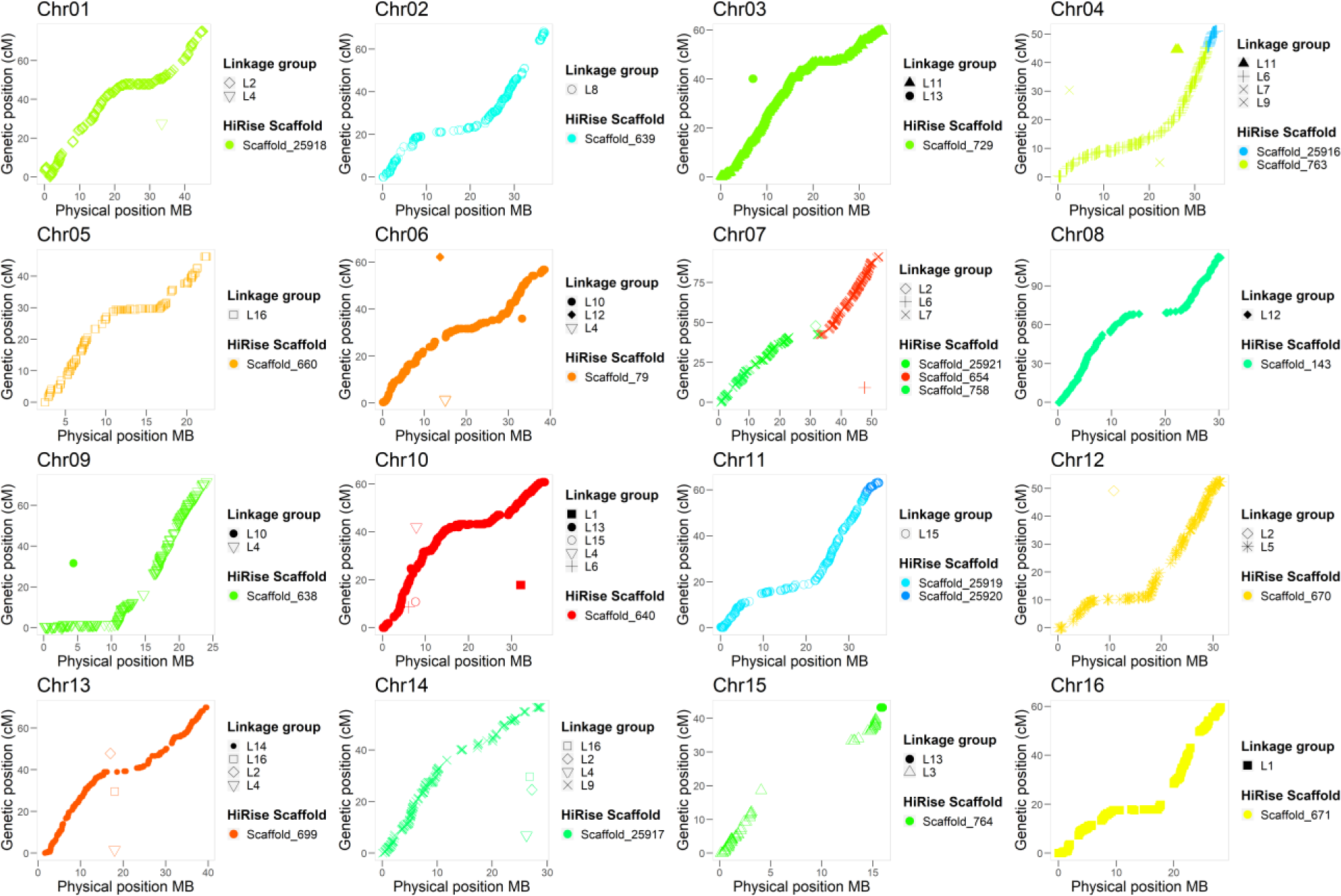
Collinearity between the high-density ‘Chandler’ genetic map of (Marrano et al. 2019) and the 16 chromosomal pseudomolecules of Chandler v2.0.

Second, we anchored the HiRise assembly to the Chandler genetic map used by (Luo et al. 2015) to construct a walnut physical map. In total, 972 of the mapped markers (1,525 SNPs) aligned uniquely on the same 35 HiRise scaffolds anchored to the linkage map mentioned above. Overall, we observed almost perfect collinearity between the HiRise assembly and both Chandler genetic maps (**Figure 1, Supplemental Figure S3**). Therefore, we oriented, ordered, and named the HiRise scaffolds consistent with the linkage map of (Luo et al. 2015), generating the final 16 chromosomal pseudomolecules of *J. regia* Chandler.

These 16 contiguous chromosomal scaffolds account for 95% of the final walnut reference genome v2.0, with an N50 size of 35 Mb. Chandler v2.0 has a total length of 576,258,700 bps, of which only 20.9 Mb are fragmented in 2,631 small unanchored scaffolds (> 1 kb; **Table 1**). The larger genome size of Chandler v2.0 compared to the recently published genome assembly of the cv. Serr (JrSerr_v1.0; 534.7 Mb) (Zhu et al. 2019) can be explained by structural variation (e.g., copy number and presence/absence variants), whose central role in explaining intraspecific genomic and phenotypic diversity has been reported in different plant species (Springer et al. 2009; Marroni et al. 2014). In addition, the higher number of unanchored scaffolds in Chandler v2.0 compared to JrSerr_v1.0 can represent autozygous genomic regions of Chandler, devoid of segregating markers and, therefore, difficult to anchor to linkage genetic maps (Luo et al. 2015), as also suggested by the higher fixation index (*F*) observed in Chandler (0.03) than Serr (-0.29) in previous genetic surveys (Marrano et al. 2018).

We identified telomere sequences at both ends for nine of the chromosome scaffolds, on one end of the other seven chromosomes and one end of seven unanchored scaffolds. Also, all 16 chromosomes had centromeric repeats in the middle, alongside regions with low recombination rates (**Figure 2**).

**Figure 2.**
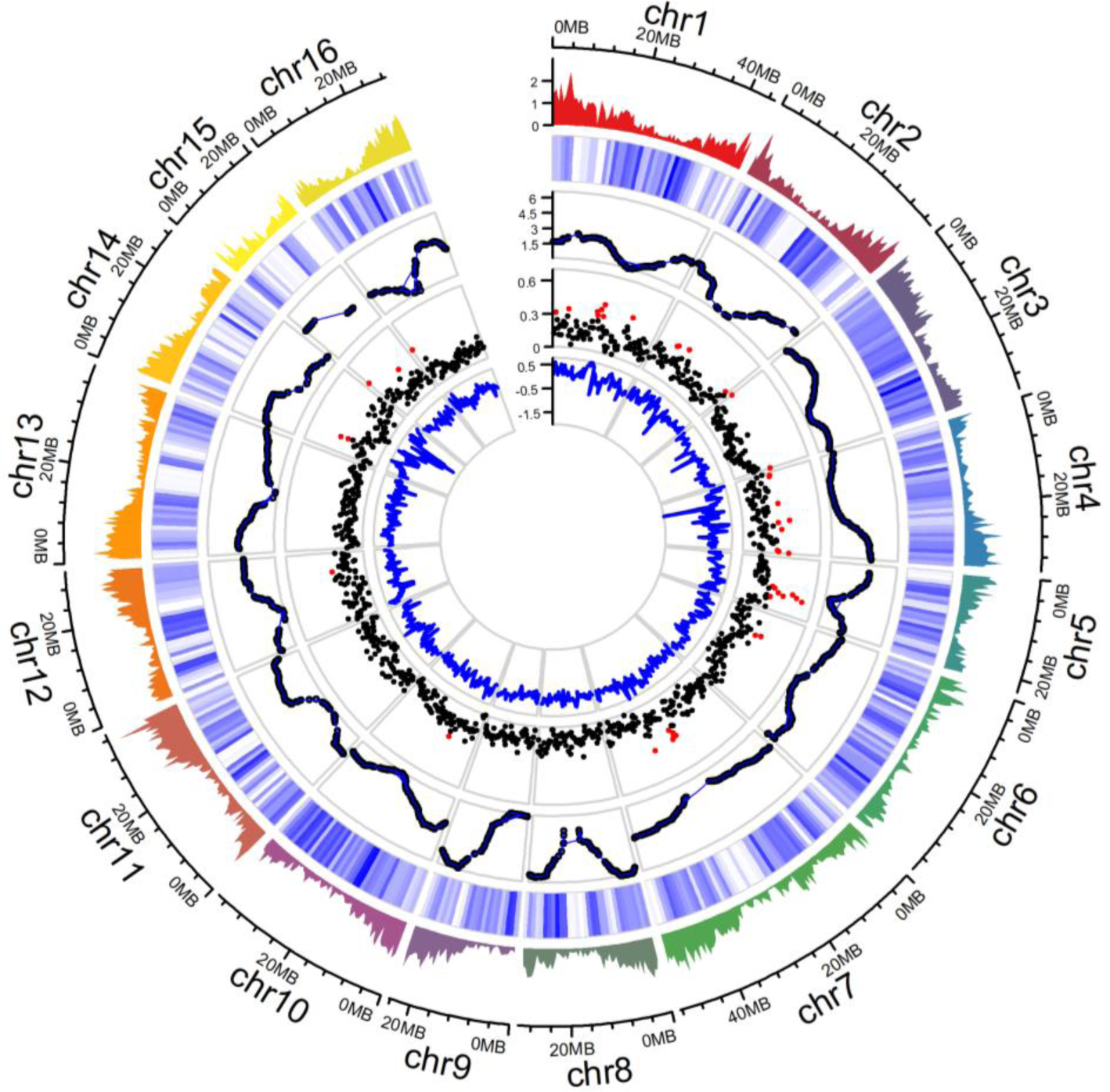
Summary of gene distribution and genetic diversity across the 16 chromosomes of Chandler v2.0. Tracks from outside to inside: (***i***) gene density of Chandler v2.0 in 1-Mb windows; (***ii***) Chandler heterozygosity in 1-Mb windows (white = low heterozygosity; blue = high heterozygosity); (***iii***) Recombination rate for sliding windows of 10 Mb (average = 2.63 cM/Mb); (***iv***) FST in 500-kb windows. Windows in the 95 percentiles of the FST distribution are highlighted in red; (***v***) ROD values for 500-kb windows.

To assess the sequence accuracy of Chandler v2.0, we first compared the scaffold sequences of Chandler v2.0 with the previous version of the walnut reference genome, generating 838,173 alignments with sequence identity averaging 94.11%. We then mapped the Illumina whole-genome shotgun data (Martínez-García et al., 2016) against the new chromosome-scale genome. The alignment resulted in 64,950,691,681 bps mapped, of which 407,450,406 were single-base mismatches, consistent with an Illumina sequence accuracy rate of 99.5%.

### Repeat annotation

Almost half (49.68%) of the new Chandler v2.0 is repetitive, similarly to the previous version of the walnut reference genome (51.19%). As in most plant genomes, interspersed repeats (45.13%) were the most abundant type of repeats, with retrotransposons at 18.55% and DNA transposons at 3.05%. *Gypsies* (6.58%) and *Copias* (4.1%) were the most represented classes of long-terminal retrotransposons (LTR), and, though widely dispersed throughout the genome, they were distributed differently along the 16 chromosomes (**Supplemental Figure S4**): the *Gypsies* LTRs were more abundant alongside the centromeres, where, instead, the density of the *Copia* LTRs decreased, consistent with (Zhu et al. 2019). L1/LINE (long-interspersed nuclear elements), which possess a poly(A) tail and two open reading frames (ORFs) for autonomous retrotransposition, was the largest class of non-LTRs at 6.98% of the genome. Simple repeats (4.29%) and low-complexity regions (0.26%) were also found.

### PacBio IsoSeq sequencing and gene annotation

A fragmented reference genome can severely hamper the accuracy of gene prediction, because many genes will be broken across multiple small contigs (false negatives), and because multiple fragments of the same gene may be annotated separately (false positives). Also, transcriptome assemblies generated using second-generation (Illumina) sequencing data are likely to miss many transcripts due to the very short read lengths (Minoche et al. 2015).

To improve the gene prediction accuracy of Chandler v2.0, we used the “Isoform Sequencing” (Iso-Seq) method, developed by Pacific Biosciences (PacBio), which can generate full-length transcripts up to 10 kb, allowing for accurate determination of exon-intron structure by alignment of the transcripts to the assembly (Rhoads and Au 2015). The high error rate of PacBio sequencing can be greatly reduced using circular consensus sequence (CCS), in which a transcript is circularized and then sequenced repeatedly to self-correct the errors. We applied PacBio IsoSeq to sequence full-length transcripts from nine tissues, chosen to cover most of the transcript diversity in walnut (**Supplemental Table S3**). Across the four SMRT cells, we obtained 26,328,087 subreads with a mean length of 1,188 bp (**Supplemental Table S4**) and CCSs ranging from 13K to 142K per library (**Supplemental Table S5**). Out of the 745,730 full-length non-chimeric (FLnc) transcripts, 68,225 were classified as high quality, FL (HQ FL) consensus transcript sequences, with an average length of 1,357 bp (**Supplemental Table S5**). Catkin 1-inch elongated (CAT1), shoot, and root yielded the lowest number of HQ FL transcripts, while pollen and leaf had the lowest number of HQ consensus clusters obtained per CCS after polishing (**Supplemental Table S5**). These results can be explained by lower cDNA quality or fewer inserts of full-length transcripts from these tissues during the cDNA pooling and library preparation. Nevertheless, more than 99% of the HQ FL transcripts aligned onto the new chromosomal-level walnut reference genome (**Supplemental Table S6**).

By combining the HQ FL transcripts with available *Juglans* transcriptome sequences, we identified 40,491 gene models, which are more than those annotated in Chandler v1.0 but fewer than the predicted genes in the NCBI RefSeq *J. regia* annotation generated with the first version of the reference genome (**Table 2**). This result suggests that the new chromosome-scale genome, along with the availability of full-length transcripts, allowed us to identify genes missed during the annotation of Chandler v1.0, as well as to remove false-positive predictions. Also, the mean gene length in Chandler v2.0 was higher than the previous gene annotations (**Table 2**), a consequence of the increased contiguity of the new chromosome-scale reference genome.

**Table 2.** Statistics on the gene annotation of Chandler v2.0 compared to the previous gene annotations of the Chandler genome.

|  | Chandler v2.0 | Chandler v1.0* | Chandler RefSeq v1.0 |
| --- | --- | --- | --- |
| Number of genes | 40,491 | 32,496 | 41,188 |
| **Average gene length (bp)** | 5,776 | 4,358 | 4,641 |
| **Single-exon transcripts** | 6,616 | 6,247 | 6,749 |
| **Average CDS length (bp)** | 1,335 | 1,222 | 1,336 |
| **Number of exons** | 244,238 | 172,273 | 230,261 |
| **Average exon length (bp)** | 257.1 | 229.5 | 314 |
| **Number of Introns** | 203,157 | 139,775 | 181,419 |
| **Average intron length** | 856.9 | 730 | 835 |

\* (Martínez-García et al. 2016)

The average gene density of Chandler v2.0 was 19.28 genes per 100 kb, with higher gene content in the proximity of telomeric regions (**Figure 2**), consistent with other plant genomes (Maccaferri et al. 2019; Linsmith et al. 2019). In addition, 92% (37,102) of the predicted gene models of Chandler v2.0 was supported by expression data, and 97% showed high similarity with a protein-coding transcript from either the *J. regia* RefSeq v1 gene set or a protein from the wider NCBI RefSeq plant database (**Supplemental Table S7**), highlighting the accuracy and robustness of the Chandler v2.0 genome annotation. Overall, Chandler v2.0 contained 339 newly predicted gene models, with a mean length of 1,533 bp. Of these new predicted gene models, 150 (44%) and 329 (97%) were supported by PacBio IsoSeq and Illumina RNA-seq data (Martínez-García et al., 2016), respectively. Thus, the failure to identify these genes in Chandler v1.0 was most likely related to its low contiguity than to the lack of the gene transcripts in the RNA-seq data.

Out of the 41,081 transcripts identified, 84% were multi-exonic, with, on average, 5.94 exons each and a mean exon length of 257.2 bp (**Table 2**). The mean number of introns per gene was 5.9, with a length ranging from 20 bp to 493 kb. These values are similar to those observed in the previous gene annotations of Chandler, except for the number of exons and introns which was higher in Chandler v2.0. Also, introns were longer, on average, contributing to the higher mean gene length observed in Chandler v2.0. The majority of intron/exon junctions were GT/AG-motif (98.2%), even though alternative splicing with non-canonical motifs was also observed (GC/AG – 0.8%; AT/AC – 0.11%). Almost 90% (36,438) of the coding sequences were full-length with canonical start and stop codons, while 4,525 presented either a start or a stop codon. This result represents a great improvement compared to Chandler v1.0, where only 48% of the predicted gene models were complete (Martínez-García et al. 2016).

Also, we observed that 568 gene models had from two to four transcript isoforms each, with a mean length of 7,080 bp. Out of the 1,158 isoforms identified, 339 were covered by FL HQ transcripts in at least one tissue, while 835 were expressed in at least one of the 20 tissues (Martínez-García et al., 2016), which most likely covered higher gene diversity compared to the nine tissues used for PacBio IsoSeq. On average, the Illumina isoforms (7,220 bp) were longer than the PacBio isoforms (6,044 bp). By running the EnTAP functional annotation pipeline with the entire NCBI RefSeq plant database (Hart et al. 2018), we observed that 672 isoforms were annotated with a plant protein, while the remaining 486 transcript isoforms were not identified in the previous walnut gene annotation.

Of the 41,103 gene models, 83% were annotated with a plant protein, and 84% had a known Pfam domain. Also, 33,034 models were annotated with 8,244 different Gene Ontology (GO) terms. The three most common biological processes were regulation of transcription (2%), defense response (1.43%), and DNA recombination (1.2%; **Supplemental Figure S5**). ATP binding (7.7%), metal ion binding (7.1%) and DNA-binding transcription factor activity (3.7%; **Supplemental Figure S6**) were the most abundant molecular functions, while nucleus (13%), integral component of membrane (10.3%) and plasma membrane (8%) were the top three cellular components (**Supplemental Figure S7**).

The majority (95%) of the 1,440 core genes in the embryophyte dataset from Benchmarking Universal Single-Copy Orthologs (BUSCO) were identified in both the new Chandler genome assembly and gene space v2.0. Also, 88% of both rosids and green sets of core gene families (coreGFs) were identified in the gene annotation, confirming the high-quality and completeness of the gene space of Chandler v2.0.

### Improved assessment of proteomes with the complete genome sequence

After confirming the importance of a chromosome-scale reference genome for the improvement of gene prediction accuracy, we studied the impact of a contiguous genome on proteomic analysis. A virtual proteome, which includes all protein sequences predicted from a reference genome, is generally used to map and assign the peptides detected in mass spectrometry (MS) to specific protein-coding genes. Therefore, a fragmented assembly of the reference genome can lead to an inaccurate prediction of a species’ proteome and, then, a miss-identification of the proteins expressed in specific tissues at particular stages (Jamet and Santoni 2018).

We analyzed the proteomic data generated from samples encompassing different developing stages of the male walnut flower (catkin) and pure pollen, by using the virtual proteomes predicted from the gene annotation of the new chromosome-scale genome and Chandler v1.0 (NCBI RefSeq). Considering all tissues analyzed, we identified fewer unique peptides (43,083) with the new chromosome-scale walnut genome than with Chandler v1.0 (44.679). Also, 6,966 unique proteins were detected with Chandler v2.0 against the 8,802 found using version 1 as a search database (**Supplemental Table S8-S9**). Most likely, the NCBI proteomic database based on the fragmented Chandler v1.0 included artifacts resulting from an overestimation of the protein-coding genes. Therefore, the new chromosome-scale genome allows accurate estimation of the proteomic changes occurring in the different vegetative and reproductive stages of walnut, which is fundamental to fully understand the molecular bases of the observed phenotypic traits.

Sample clustering according to their protein constituents and levels showed greater similarity between immature and mature catkins and a more distinct profile between senescent catkins and pure pollen (**Figure 3**).

**Figure 3.**
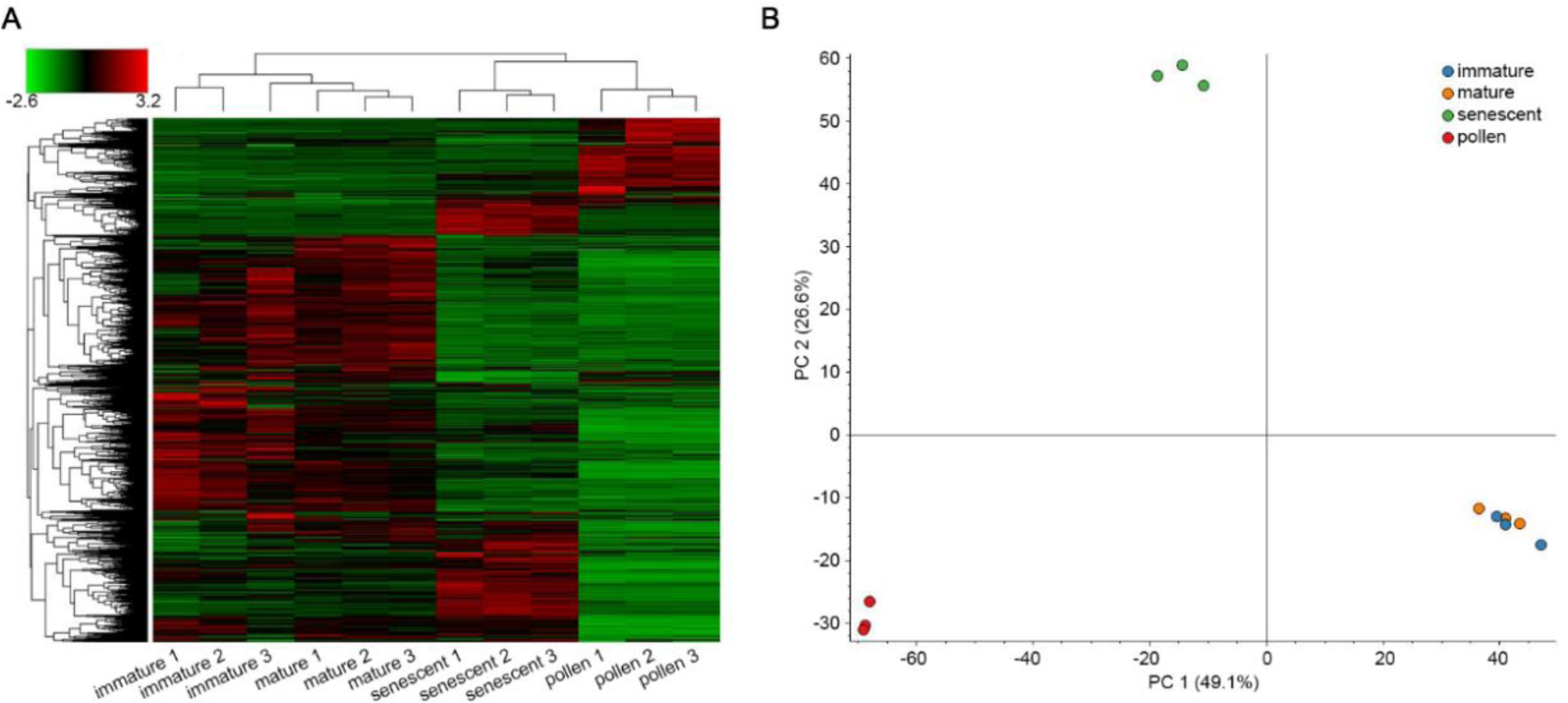
Clustering of the samples used in the proteomic analysis. (**A**) Hierarchical clustering based on Euclidian distances of normalized abundances of detected proteins. Samples are represented in columns and proteins in rows. (**B**) Principal component analysis of the 12 samples analyzed, clustering according to tissue type.

Given that ∼2% of walnut consumers have high risk of developing allergies to nuts or pollen (Costa et al. 2014), we searched the four developed proteomes for allergenic proteins listed in the WHO/IUIS Allergen Database (www.allergen.org; **Supplemental Table S10**), and additional proteins not yet registered in the allergen database but predicted in Chandler v2 as potential allergens (**Supplemental Table S9**). Four of the eight recognized allergenic proteins were detected in at least one of the catkin developmental stages, with Jug_r_5 (XP_018825777 | *Jr12_10750*) and Jug_r_7 (XP_018808763 | *Jr07_28960*) present in all sample types, including pollen (**Supplemental Table S10**). Three of the new potential allergens (**Supplemental Table S10**) are encoded by genes adjacent to known allergen-coding sequences, likely indicating gene duplications. Also, we discovered that the gene locus *Jr12_05180* encodes a non-specific lipid transfer protein (nsLTP; Jug_r_9 | XP_018813928), a potential allergen highly expressed during catkin maturation and in pollen (**Supplemental Table S10-11**). In particular, Jug_r_9 was the most abundant protein in mature and senescent catkins, and the second-most abundant in pure pollen (**Supplemental Table S10-11**). Another interesting allergen similar to Jug_r_9 (same eight cysteine configuration) is XP_018814382 | *Jr03_26970*; it decreases as the catkin matures, and is entirely absent in pollen (**Supplemental Table S10-11**). Similarly, polyphenol oxidase (PPO, XP_018858848 | *Jr03_06780*) is high in the immature catkin and almost absent in the pollen. The integration of this proteomic data with previously published transcriptomic data obtained from 20 walnut tissues (Martínez-García et al. 2016) shows high reproducibility between the methods. In both datasets, allergens Jug_r_1, 4, and 6 were not detected in catkins, while the new putative allergen Jug_r_9 was highly expressed in catkins (**Supplemental Tables S11-S12**). Also, *Jr12_05180* transcripts were not detected in any of the 20 tissues but catkin, thus confirming the strong specificity of Jug_r_9 for catkin and pollen tissue (**Supplemental Table S12**). Modeling the structure of this putative allergen reveals four predicted disulfide bonds, potentially conferring heat and protease-resistance, and further suggesting allergenic properties (**Figure 4**). Future studies will clarify the functional role of this protein and its allergenic nature.

**Figure 4.**
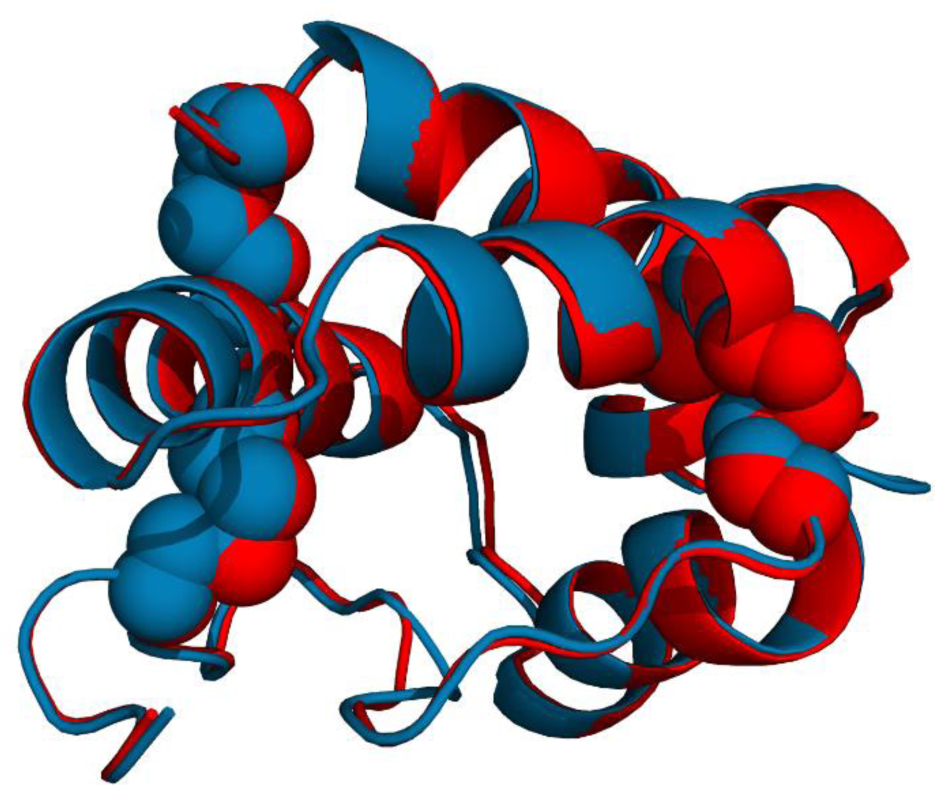
Modeled structure of the putative new allergen encoded by *Jr12_05180*. The compact structure is stabilized by four disulfide bonds, common in other allergenic proteins. The model in blue is superimposed with a homologous allergen from lentil (PDBid:2MAL) represented in red. Structure rendered with Pymol 2.3 (www.pymol.org).

The detection of new potential walnut allergens confirms the positive impact of Chandler v2.0 on proteomic studies in walnut, by providing a clearer and more precise organization of the CDSs within a genomic vicinity than the previous fragmented genome assembly v1.0.

### Chandler genomic diversity

By anchoring the HiRise assembly to the Chandler genetic map (Marrano et al. 2019), we observed highly homozygous regions in Chandler, especially on Chr15, where the genetic gap spanned 14.5 cM, corresponding to a physical distance of 9.1 Mb. A large gap on Chr15 (9.23 cM – 1.5 Mb) was also observed by (Luo et al. 2015), which suggested inbreeding as a possible cause for the lack of segregating loci in this region in Chandler, whose parents shared Payne as an ancestor. To confirm the autozygosity of Chandler on Chr15, we used the Illumina whole-genome shotgun data of Chandler and the identified polymorphisms to study its genetic diversity across the new chromosome-scale genome. We identified 2,205,835 single heterozygous polymorphisms on the 16 chromosomal pseudomolecules, with an SNP density of 4.0 SNPs per kb (**Figure 2; Supplemental Table S13**). Fifty-six 1-Mb-regions exhibited less than 377.5 SNPs (10^th^ percentile of the genome-wide SNP number distribution), and chromosomes 15, 1, 7, and 13 were the top four chromosomes in the number of low heterozygous regions (**Supplemental Table S14**). In particular, Chr15 presented nine 1-Mb windows with a significantly low number of polymorphisms, five of which span 4 Mb at the end of the chromosome. In these nine low heterozygous regions, we found 1,536 SNPs in total (**Figure 2**), of which only 25 were tiled on the Axiom *J. regia* 700K SNPs array. The absence of these polymorphisms segregating in Chandler in the SNP array could be related to either a failed identification during the SNP calling due to the highly fragmented reference genome v1.0 or with the SNP exclusion during the filtering process applied to build the genotyping array (Marrano et al. 2018). The low number of Chandler heterozygous SNPs in the array affected the end of Chr15 the most, causing a reduction in the genetic length of the corresponding linkage group (**Figure 1**), as well as leaving unexplored 4 Mb of Chandler genetic variability, which is now accessible thanks to the new chromosome-scale reference genome. The failure to anchor seven of the scaffolds with telomeric sequences can be explained by the missed detection of terminally located highly homozygous regions during genetic map constructions, due to the absence of crossing-over events with heterozygous flanking markers.

Due to the evidence of whole-genome duplication in *Juglans* genomes (Luo et al. 2015), we searched for conserved regions of synteny between Chr15 and its homologous regions in the genome, to study their level of divergence and identify other evolutionary forces as possible causes of the localized reduction of heterozygosity on Chr15. Of the 5,739 pairs of paralogous genes (8,701 genes; **Supplemental Figure S8**) identified in Chandler v2.0, 448 included genes on Chr15, and 389 of these have their respective paralogues on Chr6 (**Supplemental Figure S9**), in line with what was already reported by (Luo et al. 2015). The Chr06-Chr15 pairs of paralogous genes showed average values of divergence indexes (*KS* = 0.38; *KA* = 0.13) similar to the ones observed genome-wide for other syntelogs (*KS* = 0.4; *KA* = 0.09). Similar values of divergence were also observed for the 178 Chr06-Chr15 syntelogs (171 genes) falling within the nine low heterozygous regions on Chr15 (*KS* = 0.4, *KA* = 0.1), excluding different evolutionary rates or positive selection for these regions, and leaving inbreeding as the most reasonable explanation. Other than paralogous genes, we found 393 singletons genes in the low heterozygous regions on Chr15 of Chandler. These genes are involved in different biological processes, many of which related to signal transduction, protein phosphorylation, and response to environmental stimuli (**Supplemental Table S15**).

We further investigated the contribution of inbreeding to the high level of autozygosity on Chr 15 by visualizing the inheritance of haplotype-blocks (HB; genomic regions with little recombination) across the Chandler pedigree (**Figure 5B, Supplemental Figure S10**). We observed that Payne accounts for the entire Chandler genetic makeup (19 HBs for the total length of Chr15) inherited from Pedro (mother), where only one HB (2,08 Mb) shared the same allele of Conway-Mayette (maternal-grandfather; **Figure 5A**). Regarding the paternal genetic makeup of Chandler, 13 out of 19 HBs (9,05 Mb) on Chr15 inherited Payne alleles, providing further evidence of high inbreeding on this chromosome (**Figure 5A**). This is even more evident in assessing the number of alleles matching between Payne and Chandler across the genome: Chr15 (14 HBs for a total of 13,95 Mb; **Figure 6**) shares full allele identity with Payne for almost its entire length. Such allele matching between Chandler and its ancestor Payne also occurs on Chr1 (9 HBs for a total of 8,44 Mb), Chr4 (6 HBs - 7,68 Mb), Chr7 (21 HBs - 21,62Mb) and Chr14 (7 HBs – 12,29 Mb). These results confirm a high level of inbreeding in many genomic regions of Chandler (**Supplemental Figure S10**) and support the crucial role of Chandler v2.0 for understanding trait genetic inheritance in walnut.

**Figure 5.**
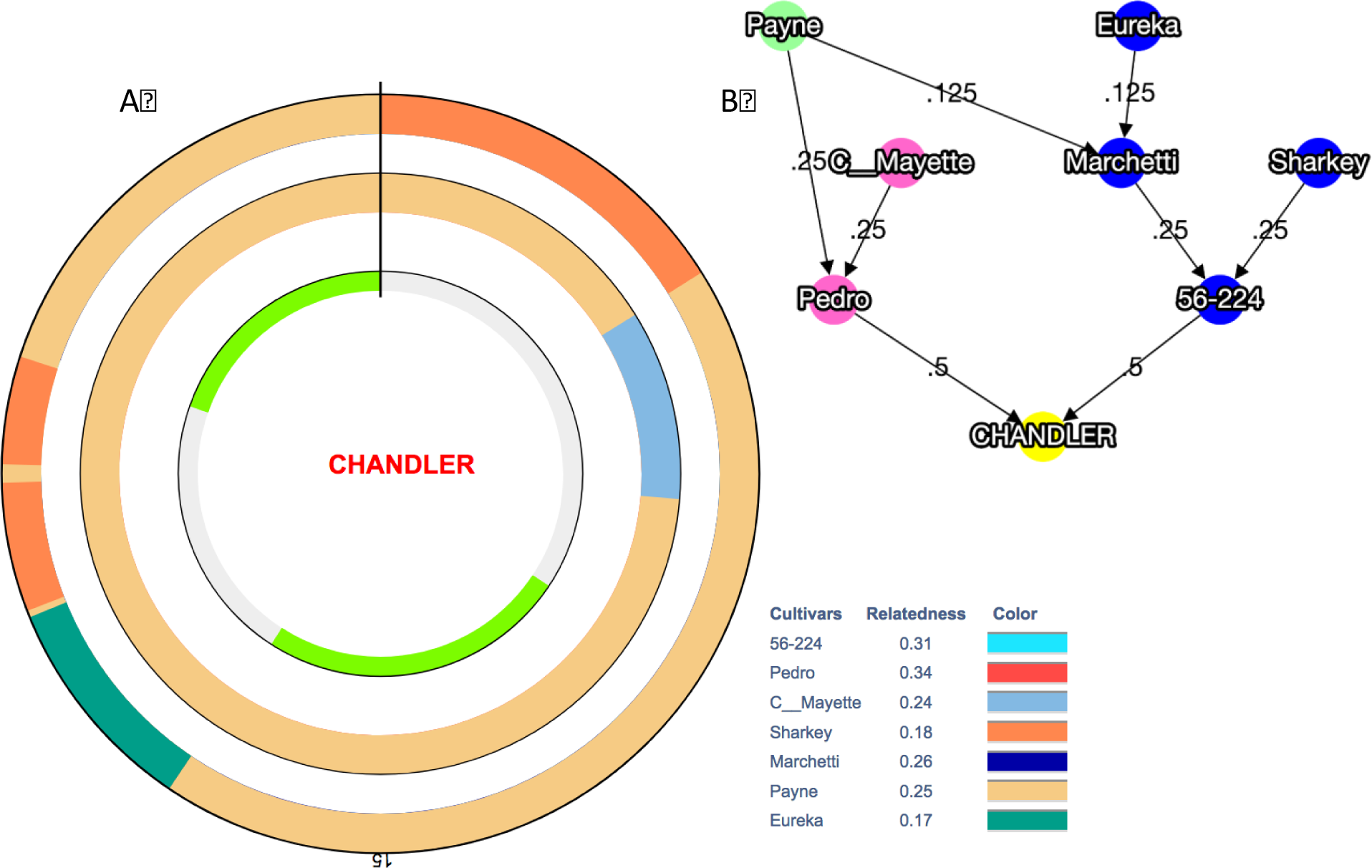
Graphical visualization of haplotype-blocks (HB) inheritance on Chr15 along with the Chandler pedigree. (**A**) The inner-circle highlights in grey two regions of heterozygosity (5 HB the first and 7 HB the second), and in light green two regions of homozygosity (3 HB the first and 4 HB the second). The circle in the middle shows maternally inherited HBs, while the HBs inherited through the paternal line are visualized in the outer circle. Payne’s haplotypes are clearly present in both parental lines. (**B**) Chandler pedigree, where Pedro is the maternal line and 56-224, the paternal line.

**Figure 6.**
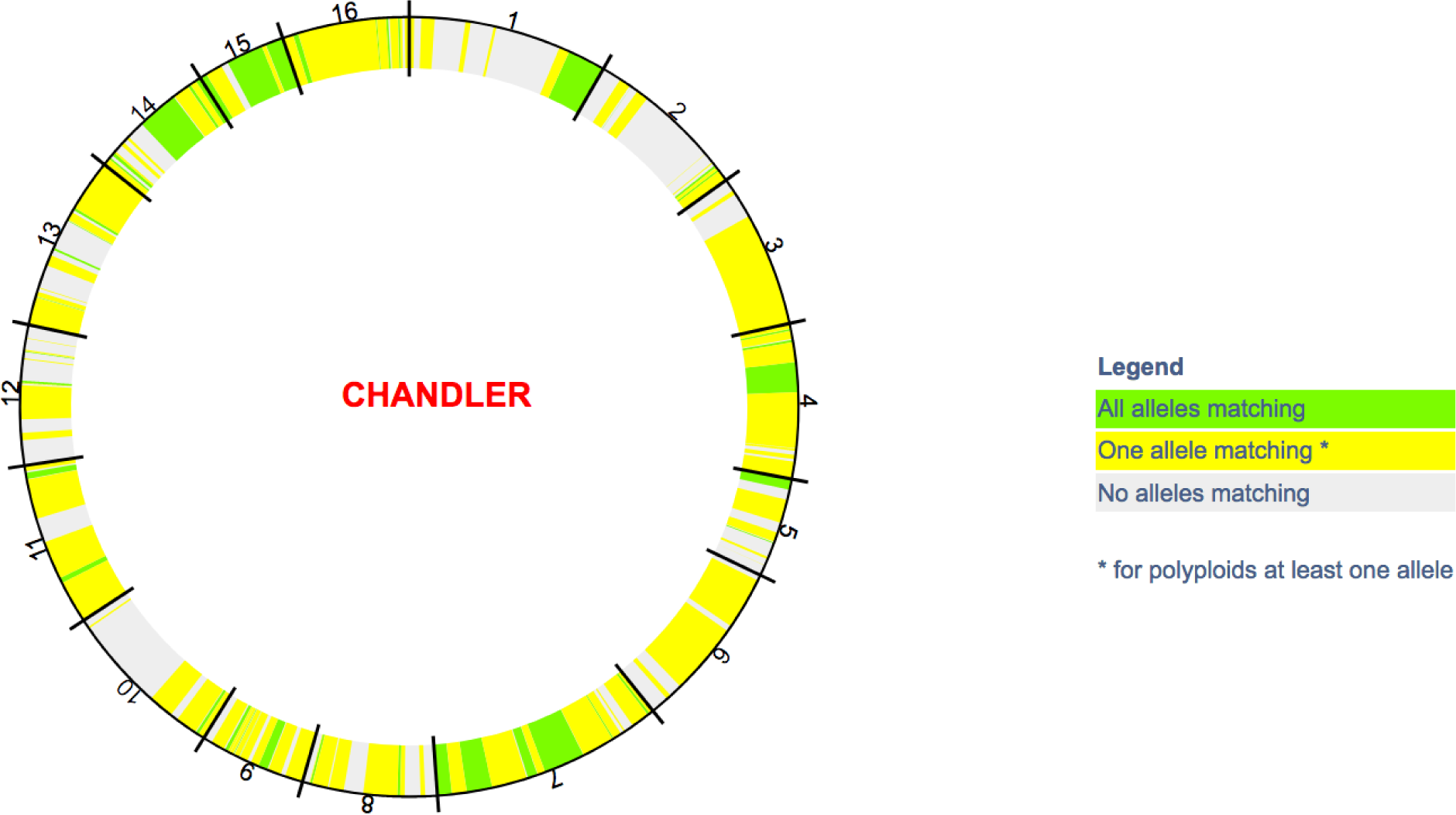
Graphical visualization of allele identity between Chandler and its ancestor Payne for all 16 chromosomes of Chandler.

### Genomic comparison between Eastern and Western walnuts

Even though numerous surveys regarding genetic diversity within walnut germplasm collections have been reported so far (Aradhya et al. 2010; Ruiz-Garcia et al. 2011), comparative analyses at the population level and genome scans for signatures of selection are still missing in Persian walnut. The availability of a chromosome-scale reference genome enables exploration of the patterns of intraspecific variation at the genomic level, providing new insight on the extraordinary phenotypic diversity present within *J. regia*.

We used the resequencing data generated for 23 founders of the Walnut Improvement Program of the University of California, Davis (UCD-WIP; **Supplemental Table S16**) (Stevens et al. 2018) to study the genome-wide genetic differentiation among walnut genotypes of different geographical provenance. We identified 14,988,422 SNPs, and over 97% of them were distributed on the 16 chromosomal pseudomolecules, with 9.4 polymorphisms per kb. A hierarchical clustering analysis (**Supplemental Figure S11**) divided the 23 founders into two major groups, including genotypes from western countries (USA, France, and Bulgaria) and Asia (China, Japan, Afghanistan), respectively, as previously reported (Marrano et al. 2018; Dangl et al. 2005). High phenotypic diversity for many traits of interest in walnut, such as phenology, nut quality, and yield, has been observed within and between germplasm collections from Western and Eastern countries (McGranahan and Leslie 1991). Walnut trees from Asia are noted for their lateral fruitfulness and precocity, rarely observed in the USA and western Europe, so that they have been used as a source of these phenotypes in different walnut breeding programs (Bernard et al. 2018b).

At a genomic level, we found a moderate differentiation (*FST* = 0.15) between Western and Eastern genotypes, except for 195 genomic windows (100 kb) that showed substantially high population differences (FST ≥ 0.36; top 5% in the whole genome). In particular, chromosomes 7, 5, 1, 4, and 2 presented about 70% of the divergent sites (**Figure 2; Supplemental Figure S12**). As suggested by the mean reduction of diversity coefficient (ROD) value (0.41), in most of the genomic regions highly differentiated, the UCD-WIP founders from the USA and Europe showed lower nucleotide diversity (π = 2.5 x 10^-4^) than the Asian genotypes (π = 5.0 x 10^-4^), consistent with (Bernard et al. 2018a) (**Figure 2; Supplemental Figure S12**). The proximity of our eastern genotypes to the supposed walnut center of domestication in Central Asia can explain the high level of diversity observed in this subgroup.

More than 60% (122) of the highly differentiated windows showed a negative value of Tajima’s D in the EU/USA subgroup (*DOcc* = -1.12), thus, suggesting that selection has been likely acting on these genomic regions in the Western genotypes (**Supplemental Figure S12**). Here we found 743 genes, with GO biological categories mostly related to signal transduction, embryo development, and response to stresses (**Supplemental Table S17**). Ten candidate selective sweeps (*DAsia* = -0.54) were also observed in the Eastern group (**Supplemental Figure S12**), which included 57 predicted genes, related to terpenoid biosynthesis, post-embryonic development, and signal transduction (**Supplemental Table S18**).

Recently, many marker-trait associations have been reported for different traits of interest in walnut, such as leafing date, nut-related phenotypes, and water use efficiency (Arab et al. 2019; Famula et al. 2019; Marrano et al. 2019). We looked to see if any of these trait-associated SNPs fell within regions highly differentiated between Western and Eastern genotypes. Three loci associated with shape index, nut roundness, and nut shape (Arab et al. 2019) are located in two genomic regions on chromosome 3 and 4 with significantly high values of FST (**Supplemental Table S19**). In both of these regions, Western genotypes presented lower genetic diversity and lower values of Tajima’s D than the Eastern walnuts. These findings may suggest that, while a selective pressure for nut shape may have occurred in the EU/USA subgroups, higher phenotypic variability can be expected for these traits in the Eastern countries. We also found that the locus AX-170770379, strongly associated with harvesting date (Marrano et al. 2019), falls within a genomic region on Chr1 with an *FST* value equal to 0.39 and lower genetic diversity in the western genotypes (ROD = 0.63; **Supplemental Table S19**). Looking at the phenotypic effect of this SNP on the harvest date of the 23 founders, we observed that most of the western genotypes are later harvesting than the eastern (**Supplemental Figure S13**), suggesting differences in the timing of phenological events between these two groups as adaptation to the different climate conditions present in their countries of origin (Gauthier and Jacobs 2011).

These results confirm the central role of a chromosome-scale genome assembly for population genetics studies, which are fundamental to study how the environment and human selection impacted walnut biology. Future resequencing projects involving larger walnut collections and covering a wider area of the global walnut distribution are necessary to confirm and interpret the observed genomic differentiation between Western and Eastern walnuts, likely helping to understand the role of this genomic divergence in the evolutionary history of Persian walnut.

## METHODS

### Oxford Nanopore sequencing and assembly

High molecular weight (HMW) DNA for Nanopore sequencing (Oxford Nanopore Technologies Inc., UK) was isolated through a nuclei extraction and lysis protocol. First, mature leaf tissue from the same tree used for the original *J. regia* genome (Martínez-García et al. 2016) was homogenized with mortar and pestle in liquid nitrogen until well ground, then added to the Nuclei Isolation Buffer (Workman et al. 2018), and stirred at 4°C for 10 minutes. The cellular homogenate was filtered through 5 layers of Miracloth (Millipore-Sigma) into a 50 mL Falcon tube, then centrifuged at 4°C for 20 minutes at 3000 x g. This speed of centrifugation was selected based on the estimated walnut genome size of 1 Gb (Zhang et al. 2012). Extracted nuclei were then lysed for 30 minutes at 65°C in the SDS-based lysis buffer described by (Mayjonade et al. 2017). Potassium acetate was added to the lysate to precipitate residual polysaccharides and proteins. The sample was incubated for 5 minutes at 4°C and then centrifuged at 4°C for 10 minutes at 2400 x g. After removing the supernatant, genomic DNA (gDNA) was ethanol precipitated, and then eluted in 10 mM Tris-Cl. Further purification of the gDNA was then performed using a Zymo Genomic DNA Clean and Concentrate column.

One µg of the isolated gDNA was prepared for sequencing using the Ligation sequencing kit (LSK108, Oxford Nanopore) following manufacturer’s protocol with an optimized end repair (100 µl sample, 14 µl enzyme, 6 µl enzyme, incubated at 20°C for 20 minutes then 65°C for 20 minutes). Libraries were sequenced for 48 hours on the Oxford Nanopore Mk1B MinION platform with the R9.4 chemistry on eight flowcells. Raw fast5 data was base-called using Albacore version 1.25.

The ONT data and Illumina reads from (Martínez-García et al. 2016) were combined using the assembly algorithm implemented in MaSuRCA v3.2.2 (Zimin et al. 2013). Super-reads were constructed using a k-mer size of 41 bp. De-duplicated scaffolds were aligned onto the previously finished *J. regia* chloroplast genome (Martínez-García et al. 2016) using “minimap2 -x asm5”, as well as to a database of 223 finished plant mitochondria (downloaded from NCBI RefSeq) using blastn with default parameters.

### Hi-C sequencing

A Hi-C library was prepared by Dovetail Genomics LLC (Santa Cruz, CA, USA) as described previously (Lieberman-Aiden et al. 2009). Briefly, for each library, chromatin was fixed in place with formaldehyde in the nucleus and then extracted. Fixed chromatin was digested with DpnII, the 5’ overhangs filled in with biotinylated nucleotides, and then free blunt ends were ligated. After ligation, crosslinks were reversed and the DNA purified from protein. Biotin that was not internal to ligated fragments was removed from the purified DNA. Purified DNA was then sheared to ∼350 bp mean fragment size. Sequencing libraries were generated using NEBNext^®^ Ultra^TM^ enzymes and Illumina-compatible adapters. Biotin-containing fragments were isolated using streptavidin beads before PCR enrichment of each library. The libraries were then sequenced on the Illumina HiSeq4000 platform.

The hybrid ONT assembly, Illumina shotgun reads (Martínez-García et al. 2016), and Dovetail Hi-C library reads were used as input data for the scaffolding software HiRise, which uses proximity ligation data to scaffold genome assemblies (Putnam et al. 2016). Shotgun and Dovetail Hi-C library sequences were aligned to the hybrid ONT assembly using a modified SNAP read mapper (http://snap.cs.berkeley.edu). The separations of Dovetail Hi-C read pairs mapped within the ONT scaffolds were analyzed by HiRise to produce a likelihood model for the genomic distance between read pairs, and the model was used to identify and break putative mis-joins, to score prospective joins, and make joins above a threshold. After scaffolding, Illumina shotgun sequences were used to close gaps between contigs, resulting in an improved HiRise assembly.

### Validation and anchoring of the HiRise assembly to Chandler genetic maps

The HiRise assembly was first anchored to the Chandler genetic map obtained by (Marrano et al. 2019) from a 312 offspring F1 population ‘Chandler x Idaho’ genotyped with the latest Axiom *J. regia* 700K SNP array. SNP probes (71-mers including the SNP site) from the Axiom *J. regia* 700K SNP array were aligned onto the HiRise assembly filtering out alignments with probe/reference identity lower than 98%, covering less than 95% of the probe length or aligning multiple times on the genome. Retained markers with a unique segregation profile were then used to anchor the HiRise scaffolds. The same procedure was also followed to anchor the HiRise assembly to the Chandler genetic map used to construct a walnut bacterial artificial chromosome (BAC) clone-based physical map by (Luo et al. 2015). The final ordering of scaffolds was performed by taking into consideration the marker genetic map position, and, in the final sequence, consecutive scaffolds were separated by sequences of 100,000 Ns.

The tandem repeat finder program (trf v4.09; (Benson 1999) was run using the recommended parameters (max mismatch delta PM PI minscore maxperiod, 2 7 7 80 10 50 500 resp.) to identify repeat elements up to 500 bp long. A histogram of repeat unit lengths was generated, and peaks at 7, 29, 33, 44, 154, and 308 bp were identified. From this data, a consensus sequence corresponding to each peak was selected. All of these repeat sequences were aligned onto the HiRise assembly using ‘nucmer’ from the MUMmer4 package (Marçais et al. 2018) with a minimum match length of 7 to capture the telomeric repeat. Based on the positions of these alignments along the chromosomes and contigs, we identified the 7-mer as the telomeric repeat and the 154-mer and 308-mer as centromeric repeats.

Recombination rate was estimated within sliding windows of 10 Mb with a step of 1 Mb along the chromosome sequence by using the high-density genetic map of Chandler (Marrano et al. 2019) and the R/MareyMap package v 1.3.4 (Rezvoy et al. 2007). To evaluate Chandler v2.0 error rate, the two assemblies, Chandler v1.0 and 2.0, were aligned to each other using the ‘nucmer’ program (Marçais et al., 2018).

### RNA preparation

Five walnut tissues (leaf, catkin 1-inch elongated; catkin 3-inches elongated, pistillate flower, and pollen) were collected from ‘Chandler’ trees at the UCD walnut orchards. Four additional samples (somatic embryo, callus, shoot, and roots) were taken from tissue culture material of ‘Chandler’. Several grams of each tissue were ground in liquid nitrogen and with insoluble polyvinylpyrrolidone (PVPP; 1% w/w). RNA was isolated using the PureLink™ Plant RNA

Reagent (Invitrogen^TM^, Carlsbad, CA) following the manufacturer’s instructions, but with an additional end wash in 1 mL of 75% Ethanol. For root tissue only, RNA isolation was performed using the MagMAX^TM^ mirVana^TM^ Total RNA Isolation Kit (Applied Biosystems^TM^, Foster City, CA) as per protocol, except for the lysis step. A different lysis buffer was created adding 100 mg of sodium metabisulfite to 10 mL of guanidine buffer (guanidine thiocyanate 4M, sodium acetate 0.2M, EDTA 25 mM, PVP-40 2.5%, pH 5.0) and 1 mL of nuclease-free water. Then, 100 mg of ground root tissue were lysed in 1 mL of the new lysis buffer using a Tissue Lyser at max frequency for 2 min. The lysate was centrifuged at 4° C for 5 min at max speed. The supernatant (500 µL) was transferred to a new tube for the following steps of RNA isolation as per protocol. RNA samples were then purified, and DNase treated using the RNeasy Plant Mini Kit (Qiagen, Hilden, Germany). The RNA quality was confirmed by running an aliquot of each sample on an Experion^TM^ Automated Electrophoresis System (Bio-Rad, Hercules, CA).

### PacBio IsoSeq sequencing

Full-length cDNA Iso-Seq template libraries for PacBio IsoSeq analysis were constructed and sequenced at the DNA Technologies & Expression Analysis Core Facility of the UC Davis Genome Center. FL double-stranded cDNA was generated from total RNA (2 µg per tissue) using the Lexogen Telo^TM^ prime Full-length cDNA Kit (Lexogen, Inc., Greenland, NH, USA). Tissue-specific cDNAs were first barcoded by PCR (16-19 cycles) using IDT barcoded primers (Integrated DNA Technologies, Inc., Coralville, Iowa), and then bead-size selected with AMPure PB beads (two different size fractions of 1X and 0.4X). The nine cDNAs were pooled in equimolar ratios and used to prepare a SMRTbell™ library using the PacBio Template Prep Kit (PacBio, Menlo Park, CA). The SMRTbell™ library was then sequenced across four Sequel v2 SMRT cells with polymerase 2.1 and chemistry 2.1 (P2.1C2.1).

PacBio raw reads were processed using the Isoseq3 v.3.0 workflow following PacBio recommendations (https://github.com/PacificBiosciences/IsoSeq3). Circular consensus sequences (CCSs) were generated using the program ‘ccs’ (https://github.com/PacificBiosciences/unanimity). The CCSs were demultiplexed and cleaned of cDNA primers using the program ‘lima’ (https://github.com/pacificbiosciences/barcoding).

Afterward, CCS clustering and polishing was performed using the program ‘isoseq3’, to generate HQ FL sequences for each of the nine tissues. FLnc and HQ clusters were aligned onto the new ‘Chandler’ assembly v2.0 with minimap2 v.2.12-r827, including the parameter ‘-ax splice’ (Li 2018).

### Repeat annotation

A genome-specific repeat database was created using the ‘basic’ mode implemented in RepeatModeler v.1.0.11 (Smit and Hubley 2008). RepeatMasker v.4.0.7 was then run to mask repeats in the walnut reference genome v.2.0 and generate a GFF file (Smit et al. 2013).

### Gene prediction and functional annotation

*J. regia* RefSeq transcripts and additional *J.regia* transcripts and protein sequences downloaded from NCBI (ftp.ncbi.nlm.nih.gov/genomes/all/GCF/001/411/555/GCF_001411555.1_wgs.5d/), along with the HQ FL IsoSeq transcripts, were used as input to the PASA pipeline v.2.3.3 (Haas et al. 2003), to assemble a genome-based transcript annotation. PASA utilizes the aligners BLAT v.35 (Kent 2002) and GMAP v.2018-07-04 (Wu and Watanabe 2005), along with TransDecoder v.5.5.0 (Haas et al. 2013), which predicts open reading frames (ORFs) as genome-based GFF coordinates. The final PASA/TransDecoder GFF3 file was post-processed to name the genes and transcripts by chromosome location consistently. Functional roles were assigned to predicted peptides using Trinotate v.3.1.1 (Grabherr et al. 2011). In particular, similarity searches were performed against several public databases (i.e., Uniprot/Swiss-Prot, NCBI NR, *Vitis_vinifera.IGGP_12x, J. regia* RefSeq) using BLAST v.2.8.1, HMMER v.3.1b2, SignalP v.4.1c, and TMHMM v.2.0c.

The completeness and quality of both genome assembly and gene annotation of Chandler v.2.0 were estimated with the BUSCO method v.3 (1,440 core genes in the embryophyte dataset) (Simão et al. 2015), and the sets of coreGFs of green plants (2,928 coreGFs) and rosids (6,092 coreGFs) from PLAZA v.2.5 (Veeckman et al. 2016). Also, RNA-Seq data generated for 20 tissues (see Martínez-García et al., 2016) were aligned to the reference genome (v1 and v2) with HISAT2 (Kim et al. 2015). The alignments of the 20 RNA-seq data and the FL transcripts along with the new genome annotation v2.0 were then used as input to StringTie v.2.0 (Pertea et al. 2016) to estimate expression levels in both fragments per kilobase per million reads (FPKM) and transcripts per million (TPM) for each transcript in the v2 annotation.

The percent identity and coverage of each *J. regia* transcript compared to proteins in the NCBI plant RefSeq database was also determined by running the EnTAP pipeline v.0.9.0 (Hart et al. 2018).

### Label-free shotgun proteomics

Plant tissues of immature, intermediate, mature catkins and pure pollen from three individual trees of Chandler at the UCD walnut orchards were collected and frozen immediately in dry ice. Tissues were then further frozen in liquid nitrogen in the laboratory and ground with mortar and pestle. Five hundred milligrams of each sample were used for total protein extraction, following the procedure for recalcitrant plant tissues of (Valerie et al. 2006), with a modification in the final buffer used to resuspend the protein pellet, consisting of 8M urea in 50mM triethylammonium bicarbonate (TEAB). One hundred micrograms of total protein from each sample were then used for proteomics.

Initially, 5 mM dithiothreitol (DTT) was added and incubated at 37°C for 30 min and 1,000 rpm shaking. Next, 15 mM iodoacetamide (IAA) was added, followed by incubation at room temperature for 30 min. The IAA was then neutralized with 30 mM DTT in incubation for 10 min. Lys-C/trypsin then was added (1:25 enzyme: total protein) followed by 4 h incubation at 37°C. After, TEAB (550 μl of 50 mM) was added to dilute the urea and activate trypsin digestion overnight. The digested peptides were desalted with Aspire RP30 Desalting Tips (Thermo Scientific), vacuum dried, and suspended in 45 μl of 50 mM TEAB. Peptides were quantified by Pierce quantitative fluorometric assay (Thermo Scientific) and 1 μg analyzed on a QExactive mass spectrometer (Thermo Scientific) coupled with an Easy-LC source (Thermo Scientific) and a nanospray ionization source. The peptides were loaded onto a Trap (100 microns, C18 100 Å 5U) and desalted online before separation using a reversed-phase (75 microns, C18 200 Å 3U) column. The duration of the peptide separation gradient was 60 min using 0.1% formic acid and 100% acetonitrile (ACN) for solvents A and B, respectively. The data were acquired using a data-dependent MS/MS method, which had a full scan range of 300-1,600 Da and a resolution of 70,000. The resolution of the MS/MS method was 17,500 and the insulation width 2 m/z with a normalized collision energy of 27. The nanospray source was operated using a spray voltage of 2.2 KV and a transfer capillary temperature heated to 250°C. Samples were analyzed at the UC Davis Proteome Core.

The raw data were analyzed using X! Tandem and viewed using the Scaffold Software v.4. (Proteome Software, Inc.). Samples were searched against UniProt databases appended with the cRAP database, which recognizes common laboratory contaminants. Reverse decoy databases were also applied to the database before the X! Tandem searches. The protein-coding sequences (CDS) annotated in Chandler v1.0 (NCBI accession PRJNA350852) and v2.0 were used as a reference for identification of proteins from the mass spectrometry data. The proteins identified were filtered in the Scaffold software based on the following criteria: 1.0% FDR (false discovery rate) at protein level (following the prophet algorithm: http://proteinprophet.sourceforge.net/), the minimum number of 2 peptides and 0.1% FDR at the peptide level. Structure of the walnut allergen (Jug r 9) was modelled using SWISS-MODEL (Arnold et al. 2006) based on the structure of a homologous allergen from lentil (PDBid:2MAL). Structures were superimposed using MUSTANG (2MAL:in red, walnut in blue) (Konagurthu et al. 2006).

### Chandler genomic diversity

Illumina whole-genome shotgun data of Chandler were aligned on the Chandler v2.0 with BWA (Li and Durbin 2009) with standard parameters. SNP calling was performed using SAMtools v1.9 (Li et al. 2009) and BCFtools v.2.1 (Narasimhan et al. 2016). SNP density for windows of 1 Mb was estimated using the command ‘SNPdensity’ implemented in VCFtools v0.1.16 (Danecek et al. 2011). Self-collinearity analysis to detect duplicated regions in Chandler v2.0 was performed with MCScanX (Wang et al. 2012), using a simplified GFF file of the new gene annotation and a self-BLASTP as input. To improve the power of collinearity detection, tandem duplications were excluded after running the function ‘detect_collinear_tandem_arrays’ implemented in MCScanX. Synonymous (*KS*) and nonsynonymous (*KA*) changes for syntenic protein-coding gene pairs were measured using the Perl script “add_ka_and_ks_to_collinearity.pl” implemented in MCScanX.

To explore the inbreeding level across the 16 chromosomal pseudomolecules of Chandler, haplotypes were built for 55 individuals of the UCD-WIP, including 25 founders and several commercially relevant walnut cultivars (e.g., Chandler, Howard, Tulare, Vina, Franquette) along with their parents and progenitors. All individuals were genotyped using the latest Axiom^TM^ *J. regia* 700K SNP array as described in (Marrano et al. 2018). To define SNP HBs, 26,544 unique and robust SNPs were selected and ordered according to the Chandler genome v2.0 physical map. Subsequently, for each SNP markers and individual, phasing and identification of closely linked groups of SNPs, without recombination in most of the pedigree, was performed using the software FlexQTL^TM^ (Bink et al. 2014) and PediHaplotyper (Voorrips et al. 2016) following the approach described in (Vanderzande et al. 2019) and (Voorrips et al. 2016). In particular, HB were defined by recombination sites detected in ancestral generation of Chandler.

### Genomic comparison between Eastern and Western walnuts

The resequencing data of 23 founders of the UCD-WIP (**Supplemental Table S16**)(Stevens et al. 2018) were mapped onto the Chandler v2.0 with BWA, and SNPs were called following the same procedure described above for Chandler. SNPs with no missing data and minor allele frequency (MAF) higher than 10% were retained for the following genetic analyses (7,269,224 SNPs out of the 14,988,422 identified). Hierarchical cluster analysis on a dissimilarity matrix of the 23 UCD-WIP founders was performed using R/SNPRelate v.1.18.0 (Zheng et al. 2012). Fixation index (FST) was measured between genotypes from EU/USA and Asia with VCFtools v0.1.16, setting windows of 100kb and 500kb. Genomic windows with the top 5% of FST values were selected as candidate regions for further analysis. The empirical cutoff with a low false discovery rate (5%) was verified by performing whole-genome permutation test (1000) with a custom Python script. Nucleotide diversity (π) and Tajima’s D (Tajima 1989) were also computed along the whole genome in 100-kb and 500-kb windows using VCFtools. Reduction of diversity coefficient (ROD) was estimated as 1 – (π Occ/ πAsia). The new walnut gene annotation v.2.0 was used to identify predicted genes in the candidate regions under selection. The distribution of the identified genes into different biological processes was evaluated using the weight01 method provided by the R/topGO (Alexa 2015). The Kolmogorov–Smirnov-like test was performed to assess the significance of over-representation of GO categories compared with all genes in the walnut gene prediction. Plots were obtained using the R/circlize v.0.4.6 and R/ggplot2 v.3.5.3 packages.

## DATA ACCESS

All raw and processed sequencing data generated in this study have been submitted to the NCBI BioProject database (https://www.ncbi.nlm.nih.gov/bioproject/) under accession number PRJNA291087. All SNP data have been submitted to Hardwood Genomics (https://hardwoodgenomics.org/Genome-assembly/2539069).

## ACKNOWLEDGMENTS

We thank the Californian Walnut Board for funding this project. We are also grateful to Sriema Walawage for assistance with RNA extraction, and Brett Phinney for preparing the raw proteome data.

## AUTHOR CONTRIBUTION

DBN and AM conceived and coordinated the research. REW and WT performed the HMW DNA extraction and Nanopore sequencing. AVZ, DP and SLS assembled the hybrid Illumina-ONT assembly. LB, MT, DP and SLS validated and anchored the HiRise assembly to the genetic maps. AM and BJA collected and extracted all RNA samples. MB analyzed the PacBio IsoSeq results and performed the repeat and gene annotation. AD conceived the design of the proteomic analyses; PAZ and SC generated and analyzed the proteomic data. LB called the SNPs in Chandler and the 23 UCD WIP founders, while AM carried out the analyses on walnut genomic diversity. EAD, LB and MT built and analyzed the SNP haplotypes. CAL provided all the plant material. AM wrote the manuscript, which has been revised by all coauthors.

