## Supplemental Material for "High-quality chromosome-scale assembly of the walnut (*Juglans regia* L) reference genome"

**Table S1.** Statistics on k-unitgs, super-reads and mega-reads obtained with the MaSuRCA assembler on ONT and Illumina reads.

| | Number | Average length (bp) | N50 (bp) | Coverage |
|---|---|---|---|---|
| **K-unitigs** | 11,818,995 | 170 | 160 | 3.25 |
| **Super-reads** | 3,679,106 | 777 | 982 | 4.61 |
| **Mega-reads** | 3,160,807 | 4,716.12 | 8,158 | 24.04 |

**Table S2.** Characteristics of the Chandler ON assembly.

| | Number | Total Size (bp) | N50 (bp) |
|---|---|---|---|
| ON Scaffolds | 1,498 | 560,575,767 | 1,663,083 |
| ON Contigs | 258 | 1,592,429 | 15,764 |
| Scaffolds “Chandler” v1.0 | 25,007 | 11,648,497 | 731 |
| **Total (nuclear genome)** | **26,763** | **573,816,693** | **1,609,923** |

**Table S3.** List of tissues used for PacBio IsoSeq.

| Tissue | Developmental Stage | Source | Abbreviation |
|---|---|---|---|
| Callus | - | In vitro | CALS |
| Catkin (1-inch) | Vegetative | Tree | CAT1 |
| Catkin (3-inches) | Vegetative | Tree | CAT3 |
| Leaf | Vegetative | Tree | LEAF |
| Pistillate Flower | Vegetative | Tree | PISF |
| Pollen | Vegetative | Tree | POLN |
| Root | Vegetative | Pot | ROOT |
| Shoot | Vegetative | In vitro | SHOOT |
| Somatic Embryo | Immature | In vitro | SOME |

**Table S4.** Statistics on the PacBio IsoSeq sequencing per flow-cell.

| SMRT Cell | First | Second | Third | Fourth | Total |
|---|---|---|---|---|---|
| Subreads | 12,615,171 | 4,762,871 | 3,856,461 | 5,093,584 | 26,328,087 |

| | | | | | |
|---|---|---|---|---|---|
| Subreads Total Length | 15,022,521,907 | 6,207,229,047 | 4,290,409,550 | 5,756,344,948 | 31,276,505,452 |
| Average Subread Length | 1,191 | 1,303 | 1,113 | 1,130 | 1,188 |
| Used for CCS | 12,367,203 | 4,702,605 | 3,797,625 | 5,013,184 | 25,880,617 |
| CCS generated | 488,157 | 174,534 | 134,471 | 174,331 | 971,493 |
| CCS Total Length | 698,640,953 | 280,847,348 | 169,362,257 | 227,109,058 | 1,375,959,616 |
| Average CCS Length | 1,431 | 1,609 | 1,259 | 1,303 | 1,416 |

**Table S5.** Statistics on CCSs, FLnc and HQ FL transcripts obtained per tissue with PacBio IsoSeq.

| Tissue | N. of CCSs* | Total Length (CCS) | Average Length (CCS) | N. of FLnc | Total Length (FLnc) | Average Length (FLnc) | HQ Clusters | Total Length (HQ FL) | Average Length (HQ FL) |
|---|---|---|---|---|---|---|---|---|---|
| CALS | 129,653 | 163,156,475 | 1,258 | 129,535 | 160,363,808 | 1,238 | 11,828 | 15,284,033 | 1,292 |
| CAT1 | 13,631 | 18,596,784 | 1,364 | 13,613 | 18,287,125 | 1,343 | 1,512 | 1,987,615 | 1,315 |
| CAT3 | 125,219 | 161,569,359 | 1,290 | 125,083 | 158,780,204 | 1,269 | 11,739 | 15,170,585 | 1,292 |
| LEAF | 142,314 | 195,702,629 | 1,375 | 142,168 | 192,587,202 | 1,355 | 11,940 | 16,713,706 | 1,400 |
| PISF | 108,068 | 145,732,504 | 1,349 | 107,989 | 143,491,487 | 1,329 | 10,501 | 14,204,376 | 1,353 |
| POLN | 74,310 | 108,230,041 | 1,456 | 74,179 | 106,535,501 | 1,436 | 4,573 | 6,685,431 | 1,462 |
| ROOT | 17,897 | 23,874,571 | 1,334 | 17,875 | 23,480,651 | 1,314 | 2,252 | 2,852,647 | 1,267 |
| SHOOT | 14,508 | 19,327,322 | 1,332 | 14,494 | 19,003,828 | 1,311 | 1,400 | 1,743,863 | 1,246 |
| SOME | 120,935 | 187,564,762 | 1,551 | 120,794 | 184,845,798 | 1,530 | 12,480 | 19,783,294 | 1,585 |
| Total | 746,535 | 1,023,754,447 | 1,368 | 745,730 | 1,007,375,604 | 1,347 | 68,225 | 94,425,550 | 1,357 |

***above threshold**

**Table S6.** Percentage of FLnc and HQ FL transcripts aligned on the new assembly per tissue.

| Tissue | FLnc mapped on Chandler v2.0 | HQ clusters mapped on Chandler v2.0 |
|---|---|---|
| CALS | 99.88% | 100.00% |
| CAT1 | 99.76% | 100.00% |
| CAT3 | 99.77% | 99.96% |
| LEAF | 99.90% | 99.99% |
| PISF | 99.89% | 99.99% |
| POLN | 97.84% | 94.18% |
| ROOT | 98.69% | 99.88% |
| SHOOT | 99.86% | 100.00% |
| SOME | 99.90% | 99.99% |

**Table S7.** Validation of gene annotation with expression data and public plant protein databases.

| | v2 transcripts (41,081) | v2 gene models (40,491) |
|---|---|---|
| supported by RNA-seq data from (Martínez-García et al., 2016) | 37,370 | 37,103 |
| Supported by PacBio IsoSeq data | 19,141 | 19,005 |
| With a hit to a *J. regia* RefSeq v1 accessions | 40,738 | 40,152 |
| Annotate with a plant protein by EnTAP using the entire NCBI RefSeq plant database | 39,868 | 39,713 |

**Table S13.** Number of SNPs and SNP density in 100-kb windows per chromosome in Chandler v2.0.

| Chr | Length | Total Number of SNPs | Mean Number of SNPs/100Kb |
|---|---|---|---|
| Chr01 | 45,207,397 | 173,403 | 382.79 |
| Chr02 | 37,821,870 | 145,689 | 385.42 |
| Chr03 | 35,064,427 | 194,559 | 554.30 |
| Chr04 | 34,823,025 | 151,361 | 433.70 |
| Chr05 | 22,562,875 | 92,795 | 412.42 |

| | | | |
|---|---|---|---|
| Chr06 | 39,020,271 | 189,076 | 483.57 |
| Chr07 | 52,418,484 | 195,627 | 373.33 |
| Chr08 | 30,564,197 | 128,621 | 420.33 |
| Chr09 | 24,263,475 | 97,202 | 400.01 |
| Chr10 | 37,707,155 | 216,161 | 571.85 |
| Chr11 | 37,114,715 | 140,513 | 377.72 |
| Chr12 | 31,492,331 | 120,082 | 381.21 |
| Chr13 | 39,757,759 | 127,886 | 321.32 |
| Chr14 | 28,841,373 | 77,099 | 266.78 |
| Chr15 | 20,407,330 | 40,447 | 198.27 |
| Chr16 | 28,711,772 | 115,314 | 400.40 |
| **Tot.** | **545,778,456** | **2,205,835** | **6,363** |

**Table S14.** Regions (1 Mb) with less than 377.5 SNPs ($10^{th}$ percentile of the SNP number distribution) in Chandler.

| Chr | Window start | Window end | SNP count | Variants/kb |
|---|---|---|---|---|
| Chr01 | 5000000 | 6000000 | 11 | 0.011 |
| Chr01 | 6000000 | 7000000 | 145 | 0.145 |
| Chr01 | 11000000 | 12000000 | 99 | 0.099 |
| Chr01 | 12000000 | 13000000 | 153 | 0.153 |
| Chr01 | 37000000 | 38000000 | 278 | 0.278 |
| Chr01 | 38000000 | 39000000 | 75 | 0.075 |
| Chr01 | 43000000 | 44000000 | 163 | 0.163 |
| Chr01 | 45000000 | 46000000 | 334 | 0.334 |
| Chr02 | 14000000 | 15000000 | 184 | 0.184 |
| Chr02 | 16000000 | 17000000 | 357 | 0.357 |
| Chr02 | 18000000 | 19000000 | 203 | 0.203 |
| Chr02 | 33000000 | 34000000 | 30 | 0.03 |
| Chr02 | 34000000 | 35000000 | 83 | 0.083 |
| Chr03 | 35000000 | 36000000 | 256 | 0.256 |
| Chr05 | 1000000 | 2000000 | 123 | 0.123 |
| Chr06 | 10000000 | 11000000 | 174 | 0.174 |
| Chr06 | 39000000 | 40000000 | 3 | 0.003 |
| Chr07 | 10000000 | 11000000 | 227 | 0.227 |
| Chr07 | 26000000 | 27000000 | 112 | 0.112 |
| Chr07 | 27000000 | 28000000 | 240 | 0.24 |

| | | | | |
|---|---|---|---|---|
| Chr07 | 29000000 | 30000000 | 181 | 0.181 |
| Chr07 | 30000000 | 31000000 | 119 | 0.119 |
| Chr07 | 51000000 | 52000000 | 92 | 0.092 |
| Chr08 | 16000000 | 17000000 | 116 | 0.116 |
| Chr08 | 18000000 | 19000000 | 293 | 0.293 |
| Chr08 | 19000000 | 20000000 | 359 | 0.359 |
| Chr09 | 15000000 | 16000000 | 110 | 0.11 |
| Chr09 | 24000000 | 25000000 | 65 | 0.065 |
| Chr11 | 8000000 | 9000000 | 109 | 0.109 |
| Chr11 | 18000000 | 19000000 | 173 | 0.173 |
| Chr11 | 20000000 | 21000000 | 109 | 0.109 |
| Chr11 | 21000000 | 22000000 | 191 | 0.191 |
| Chr11 | 37000000 | 38000000 | 326 | 0.326 |
| Chr12 | 1000000 | 2000000 | 171 | 0.171 |
| Chr12 | 20000000 | 21000000 | 73 | 0.073 |
| Chr12 | 21000000 | 22000000 | 79 | 0.079 |
| Chr13 | 0 | 1000000 | 244 | 0.244 |
| Chr13 | 17000000 | 18000000 | 352 | 0.352 |
| Chr13 | 19000000 | 20000000 | 261 | 0.261 |
| Chr13 | 20000000 | 21000000 | 376 | 0.376 |
| Chr13 | 21000000 | 22000000 | 189 | 0.189 |
| Chr13 | 27000000 | 28000000 | 227 | 0.227 |
| Chr14 | 12000000 | 13000000 | 219 | 0.219 |
| Chr14 | 13000000 | 14000000 | 213 | 0.213 |
| Chr14 | 16000000 | 17000000 | 358 | 0.358 |
| Chr15 | 4000000 | 5000000 | 247 | 0.247 |
| Chr15 | 6000000 | 7000000 | 91 | 0.091 |
| Chr15 | 9000000 | 10000000 | 190 | 0.19 |
| Chr15 | 11000000 | 12000000 | 351 | 0.351 |
| Chr15 | 16000000 | 17000000 | 199 | 0.199 |
| Chr15 | 17000000 | 18000000 | 59 | 0.059 |
| Chr15 | 18000000 | 19000000 | 202 | 0.202 |
| Chr15 | 19000000 | 20000000 | 116 | 0.116 |
| Chr15 | 20000000 | 21000000 | 81 | 0.081 |
| Chr16 | 6000000 | 7000000 | 106 | 0.106 |
| Chr16 | 23000000 | 24000000 | 184 | 0.184 |

**Table S15.** Top 50 biological process GO terms for the 393 singletons genes in the low heterozygous regions on Chr15 of Chandler.

| GO.ID | Term | Total annotated genes | Number of significant genes on Chr15 | P-value |
|---|---|---|---|---|
| GO:0007165 | signal transduction | 2,682 | 49 | 9.00E-22 |
| GO:0009626 | plant-type hypersensitive response | 366 | 14 | 3.60E-19 |
| GO:0006468 | protein phosphorylation | 1,154 | 24 | 3.10E-10 |
| GO:0045088 | regulation of innate immune response | 212 | 2 | 3.10E-08 |
| GO:0051258 | protein polymerization | 105 | 2 | 4.60E-08 |
| GO:0048544 | recognition of pollen | 120 | 10 | 1.30E-06 |
| GO:0000086 | G2/M transition of mitotic cell cycle | 44 | 1 | 1.80E-06 |
| GO:0009308 | amine metabolic process | 152 | 2 | 2.40E-06 |
| GO:0072593 | reactive oxygen species metabolic process | 236 | 4 | 7.60E-06 |
| GO:0009833 | plant-type primary cell wall biogenesis | 54 | 1 | 9.50E-06 |
| GO:0009793 | embryo development ending in seed dormancy | 506 | 4 | 9.60E-06 |
| GO:0001709 | cell fate determination | 35 | 1 | 1.00E-05 |
| GO:0009267 | cellular response to starvation | 202 | 1 | 1.10E-05 |
| GO:0009636 | response to toxic substance | 306 | 4 | 1.20E-05 |
| GO:0016311 | dephosphorylation | 194 | 6 | 3.20E-05 |
| GO:0016236 | macroautophagy | 115 | 2 | 3.40E-05 |
| GO:0015074 | DNA integration | 724 | 14 | 4.00E-05 |
| GO:0009791 | post-embryonic development | 2,153 | 20 | 4.80E-05 |
| GO:0009409 | response to cold | 484 | 6 | 9.10E-05 |
| GO:0016114 | terpenoid biosynthetic process | 234 | 5 | 0.00014 |
| GO:0046777 | protein autophosphorylation | 423 | 6 | 0.0002 |
| GO:0006790 | sulfur compound metabolic process | 421 | 4 | 0.00025 |
| GO:0000272 | polysaccharide catabolic process | 294 | 7 | 0.00027 |

**Table S16.** List of 23 UCD-WIP founders used for the selective sweep analysis (Stevens et al., 2018).

| Accession Name | Origin | Abbreviation |
|---|---|---|
| UC-85-008 | China | XJG6 |
| UC-85-043-1 | Bulgaria | SH_SD |
| UC-86-011 | France/Germany | J_PUR |
| UC-91-013-5 | China | ZL5 |
| UC-91-031-8 | China | AH85 |
| UC-91-041-12 | China | AK67 |
| UC-91-056-9 | China | PRLY |

| | | |
|---|---|---|
| Conway-Mayette | USA | CMAY |
| Eureka | USA | EURK |
| Hartley | USA | HART |
| Idaho | USA | IDA |
| Lara | France | LARA |
| Manregian (PI18256) | China | MAREG |
| Marchetti | USA | MARCH |
| Meylan | France | MEYL |
| Payne | USA | PAYN |
| PI159568 | Afghanistan | PI15_8 |
| Scharsch Franquette | USA/France | S_FRA |
| Sharkey | USA | SHARK |
| Sinensis #5 | Japan | SIN5 |
| Soleze | France | SLZ |
| Waterloo | USA | WAT |
| UC-64-057 | USA | 64-057 |

**Table S17.** Top 50 biological process GO terms for the 122 windows (100 kb) with negative value of Tajima's D in the Western genotypes.

| **GO.ID** | **Term** | **Total annotated genes** | **Number of genes in the regions with negative $D_{occ}$** | **P-value** |
|---|---|---|---|---|
| GO:0007165 | signal transduction | 2,682 | 43 | 4.30E-21 |
| GO:0009626 | plant-type hypersensitive response | 366 | 3 | 5.30E-19 |
| GO:0006468 | protein phosphorylation | 1,154 | 14 | 4.90E-10 |
| GO:0051258 | protein polymerization | 105 | 7 | 4.10E-08 |
| GO:0045088 | regulation of innate immune response | 212 | 2 | 1.50E-07 |
| GO:0000086 | G2/M transition of mitotic cell cycle | 44 | 2 | 1.80E-06 |
| GO:0009308 | amine metabolic process | 152 | 5 | 3.90E-06 |
| GO:0072593 | reactive oxygen species metabolic process | 236 | 4 | 6.70E-06 |
| GO:0009636 | response to toxic substance | 306 | 5 | 8.50E-06 |
| GO:0009625 | response to insect | 28 | 1 | 1.20E-05 |
| GO:0006081 | cellular aldehyde metabolic process | 135 | 1 | 1.90E-05 |
| GO:0080027 | response to herbivore | 4 | 1 | 2.00E-05 |
| GO:0016311 | dephosphorylation | 194 | 2 | 2.50E-05 |
| GO:0009791 | post-embryonic development | 2,153 | 36 | 3.60E-05 |
| GO:0009793 | embryo development ending in seed dormancy | 506 | 8 | 3.90E-05 |
| GO:0009861 | jasmonic acid and ethylene-dependent systemic resistance | 45 | 1 | 4.60E-05 |

| | | | | |
|---|---|---|---|---|
| GO:0046777 | protein autophosphorylation | 423 | 4 | 8.80E-05 |
| GO:0015074 | DNA integration | 724 | 12 | 0.0001 |
| GO:0046839 | phospholipid dephosphorylation | 31 | 1 | 0.0001 |
| GO:0016114 | terpenoid biosynthetic process | 234 | 2 | 0.00011 |
| GO:0071555 | cell wall organization | 677 | 7 | 0.00018 |
| GO:0030244 | cellulose biosynthetic process | 120 | 2 | 0.0002 |

**Table S18.** Top 50 biological process GO terms for the 122 windows (100 kb) with negative value of Tajima's D in the Eastern genotypes.

| **GO.ID** | **Term** | **Total annotated genes** | **Number of genes in the regions with negative $D_{Asia}$** | **P-value** |
|---|---|---|---|---|
| GO:0007165 | signal transduction | 2,682 | 2 | 7.00E-22 |
| GO:0009793 | embryo development ending in seed dormancy | 506 | 1 | 9.40E-06 |
| GO:0009636 | response to toxic substance | 306 | 1 | 1.20E-05 |
| GO:0006081 | cellular aldehyde metabolic process | 135 | 2 | 2.40E-05 |
| GO:0009791 | post-embryonic development | 2,153 | 3 | 4.70E-05 |
| GO:0016114 | terpenoid biosynthetic process | 234 | 5 | 0.00014 |

**Table S19.** Marker-trait associations identified within genomic regions highly differentiated between Western and Eastern walnuts.

| Trait | SNP | Chr | Position (bp) | P-value | Window Start | Window end | N. SNPs | $F_{ST}$ | $\pi_{Asia}$ | $\pi_{Occ}$ | ROD | $D_{asia}$ | $D_{Occ}$ |
|---|---|---|---|---|---|---|---|---|---|---|---|---|---|
| Harvest Date | AX-170770379* | 1 | 6506363 | 9.2E-13 | 6500001 | 6600000 | 1526 | 0.394 | 0.006 | 0.002 | 0.629 | 1.877 | 0.771 |
| Shape index | AX-171158510** | 3 | 280565 | 1.82E-08 | 200001 | 300000 | 659 | 0.373 | 0.002 | 0.001 | 0.392 | 1.318 | 0.105 |
| Round index | AX-171158510 | 3 | 280565 | 4.38E-07 | 200001 | 300000 | 659 | 0.373 | 0.002 | 0.001 | 0.392 | 1.318 | 0.105 |
| Nut shape | AX-171105437** | 4 | 6315247 | 1.07E-05 | 6300001 | 6400000 | 2189 | 0.371 | 0.010 | 0.002 | 0.791 | 2.652 | -0.533 |
| Nut shape | AX-171105430** | 4 | 6326737 | 1.07E-05 | 6300001 | 6400000 | 2189 | 0.371 | 0.010 | 0.002 | 0.791 | 2.652 | -0.533 |

*(Marrano et al., 2019)

**(Arab et al., 2019)

**Figure S1.** Contiguity of the Chandler ON assembly and the final HiRise scaffolds. Each curve shows the fraction of the total length of the assembly present in scaffolds of a given length or smaller. The fraction of the assembly is indicated on the Y-axis and the scaffold length in base pairs is given on the X-axis. The two dashed lines mark the N50 and N90 lengths of each assembly. Scaffolds less than 1 kb are excluded.

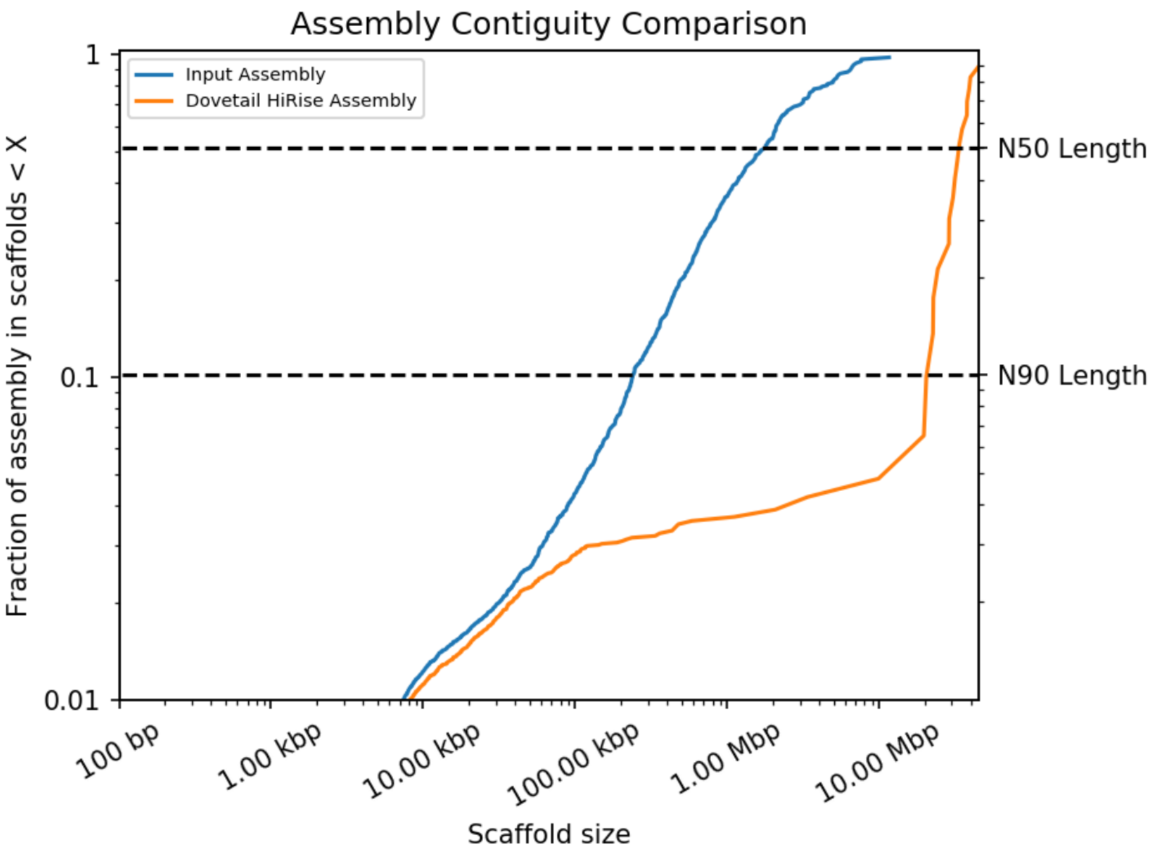

**Figure S2.** Mapping positions of the first and second read in the read pair respectively, grouped into bins. The color of each square gives the number of read pairs within that bin. White vertical and black horizontal lines show the borders between scaffolds. Scaffolds less than 1 Mb are excluded.

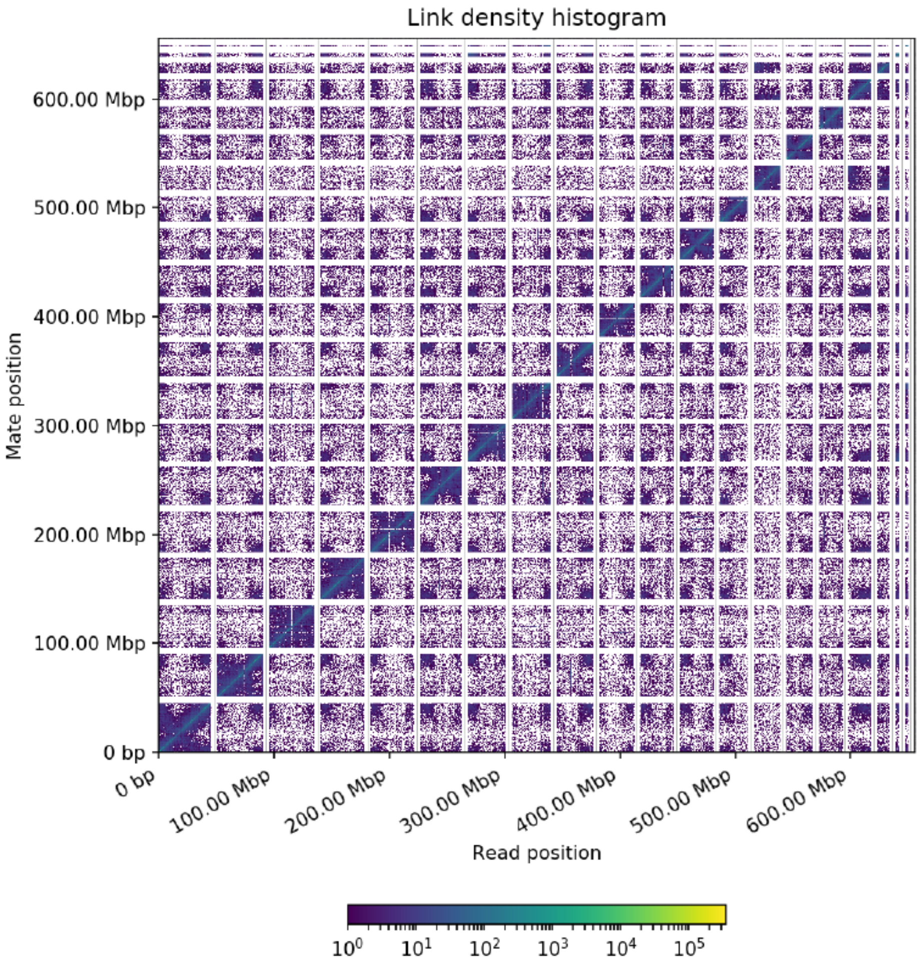

**Figure S3.** Collinearity between the 'Chandler' genetic map of (Luo et al., 2015) and the 16 chromosomal pseudomolecules of Chandler v2.0.

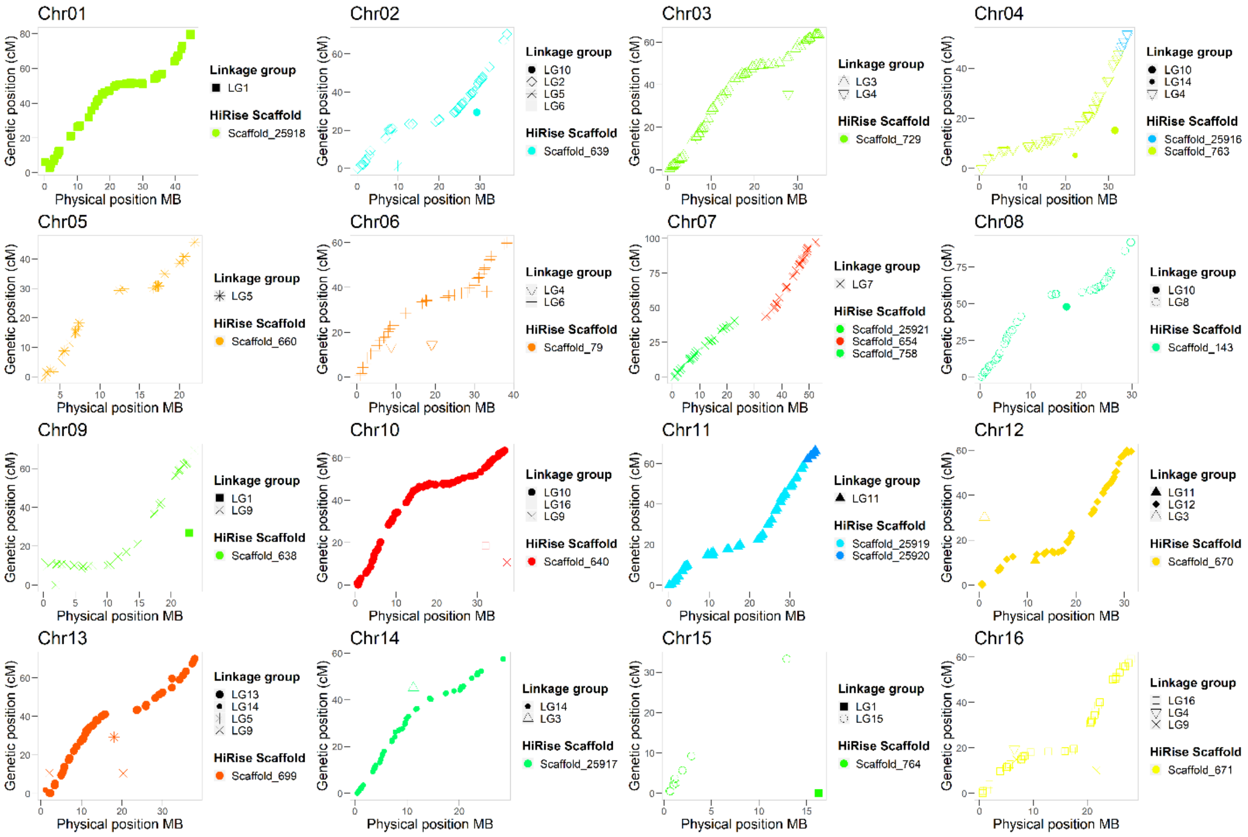

**Figure S4.** Retrotransposons distribution across the 16 chromosomes of Chandler v2.0.

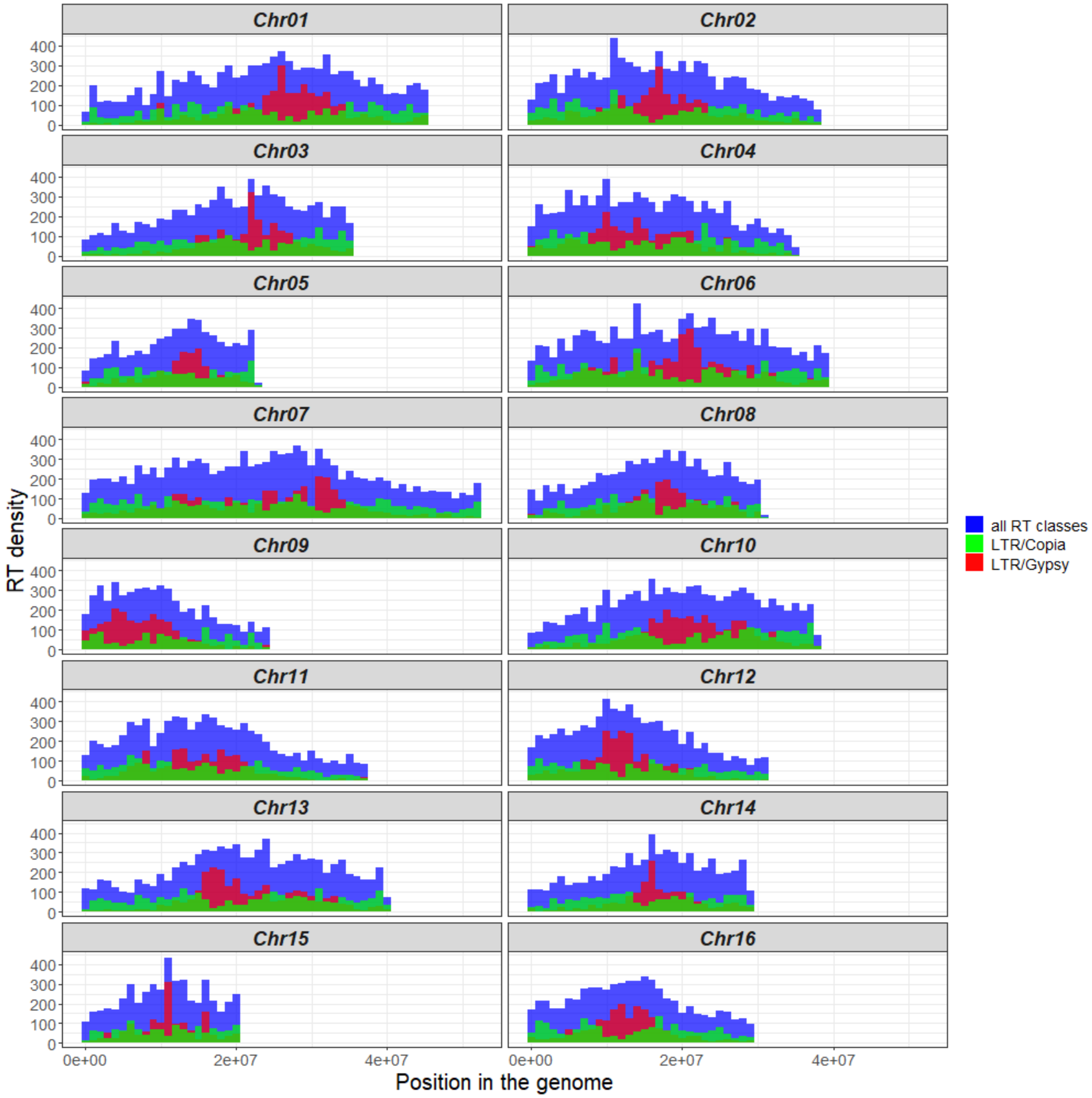

**Figure S5.** Top 20 biological process GO terms.

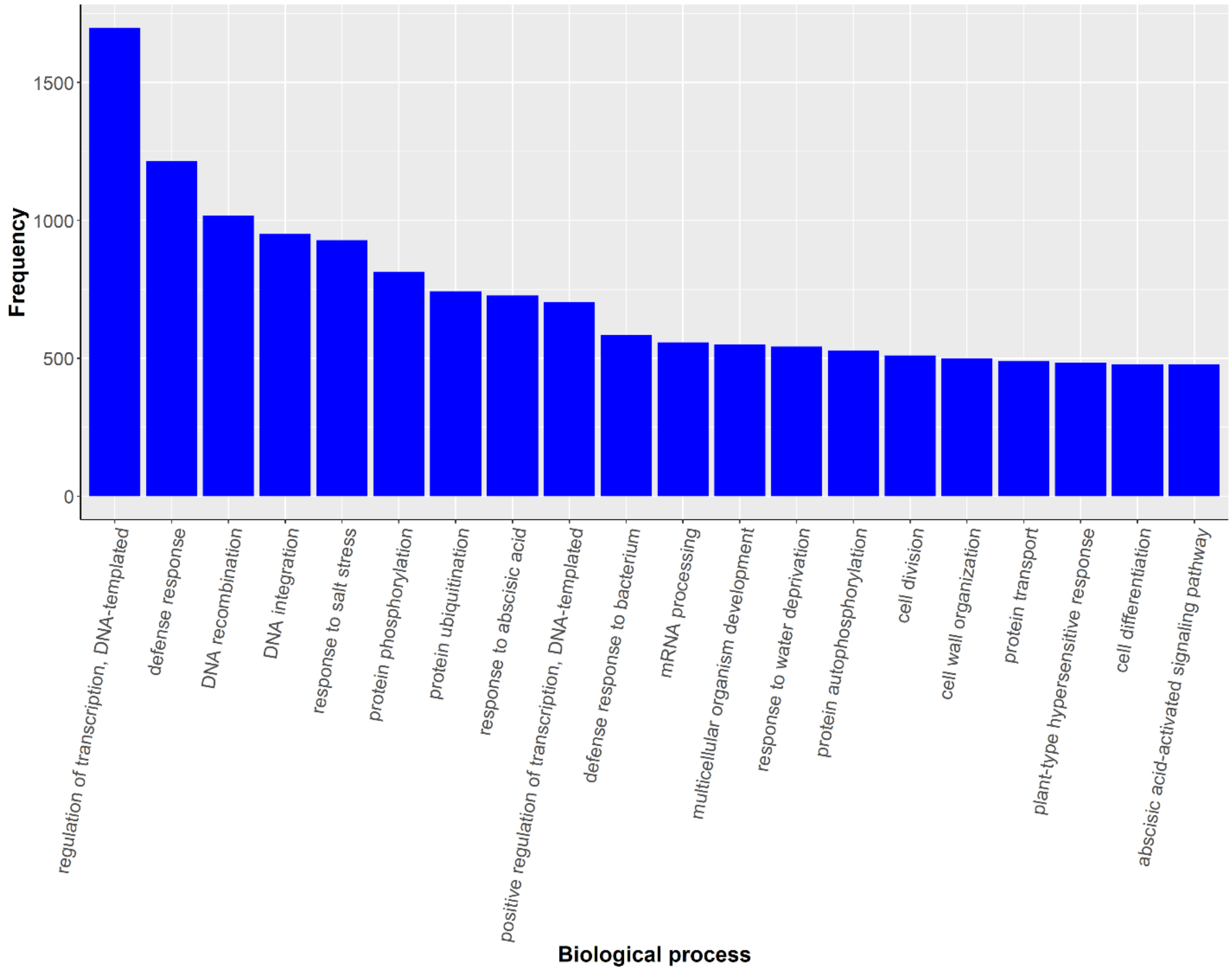

**Figure S6.** Top 20 molecular function GO terms.

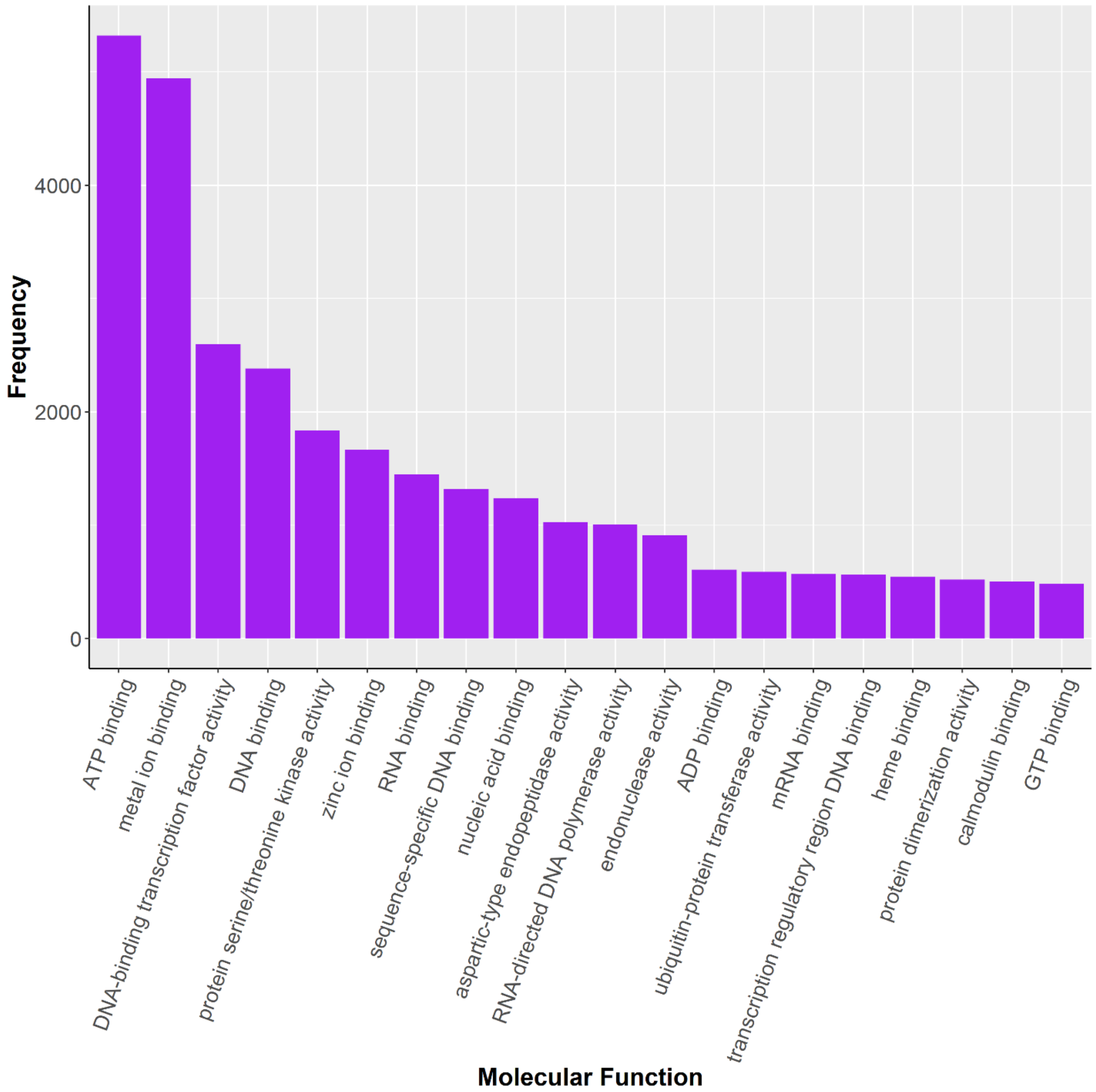

**Figure S7.** Top 20 cellular component GO terms.

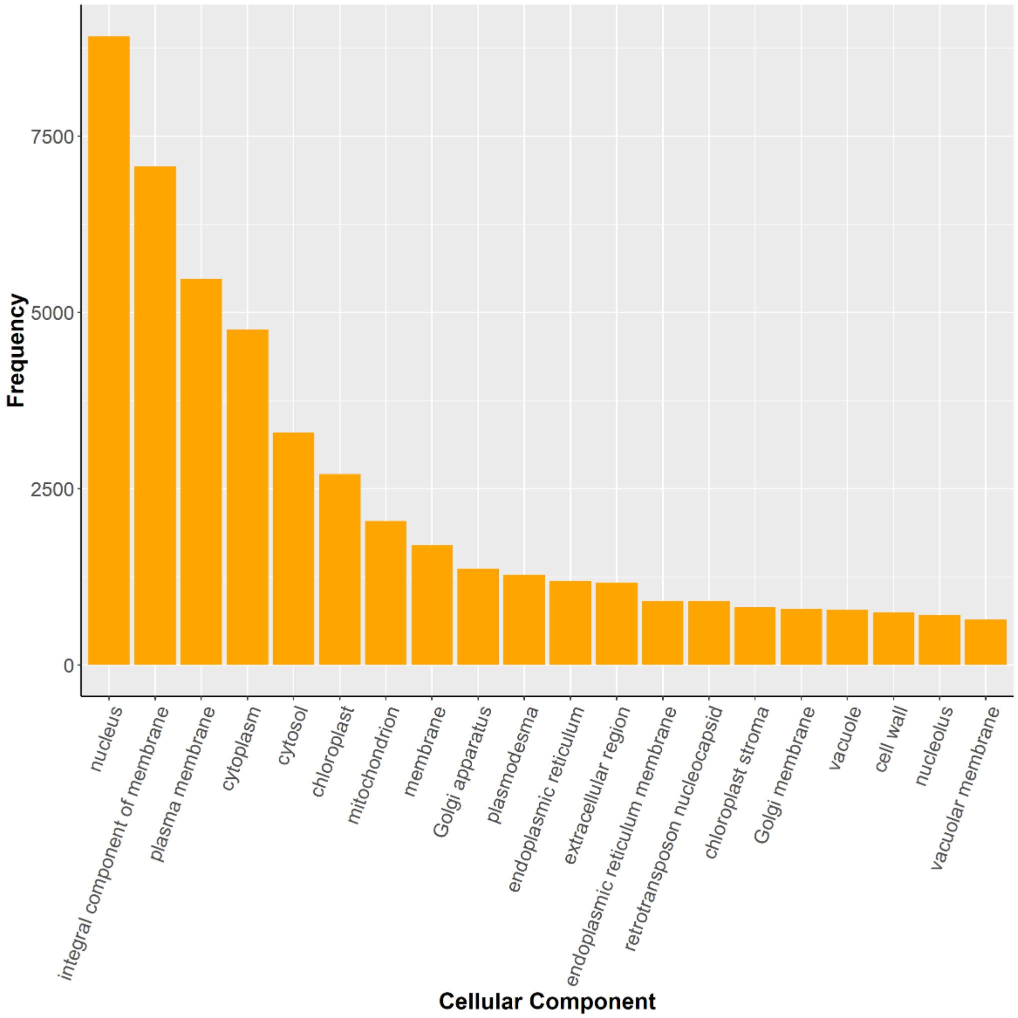

**Figure S8.** Collinear blocks among the 16 chromosomal pseudomolecules of Chandler v2.0. Different colors of dots represent different pairs of paralogous genes.

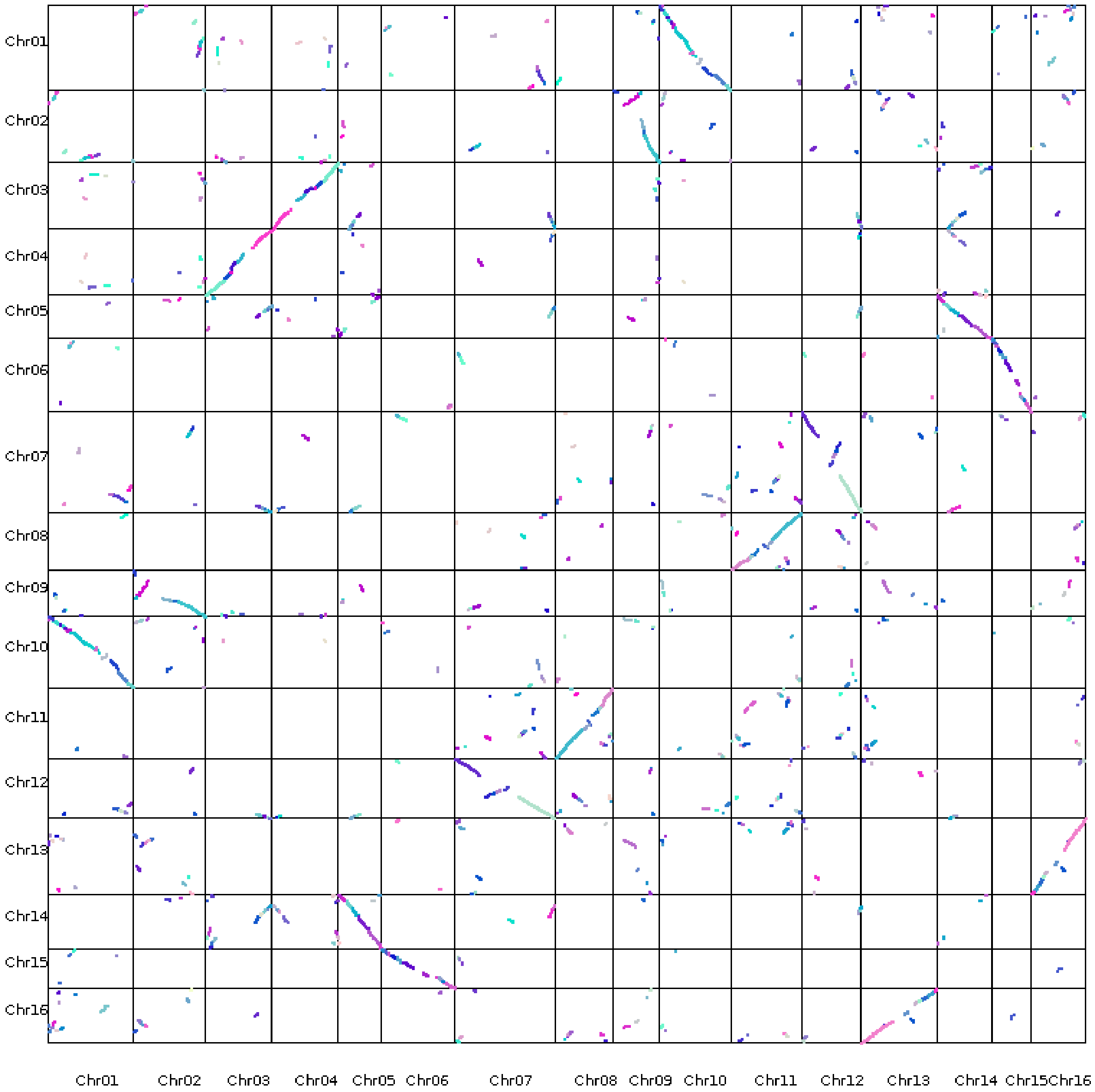

**Figure S9.** Dual synteny plot between Chr06 a- Chr15 of Chandler v2.0.

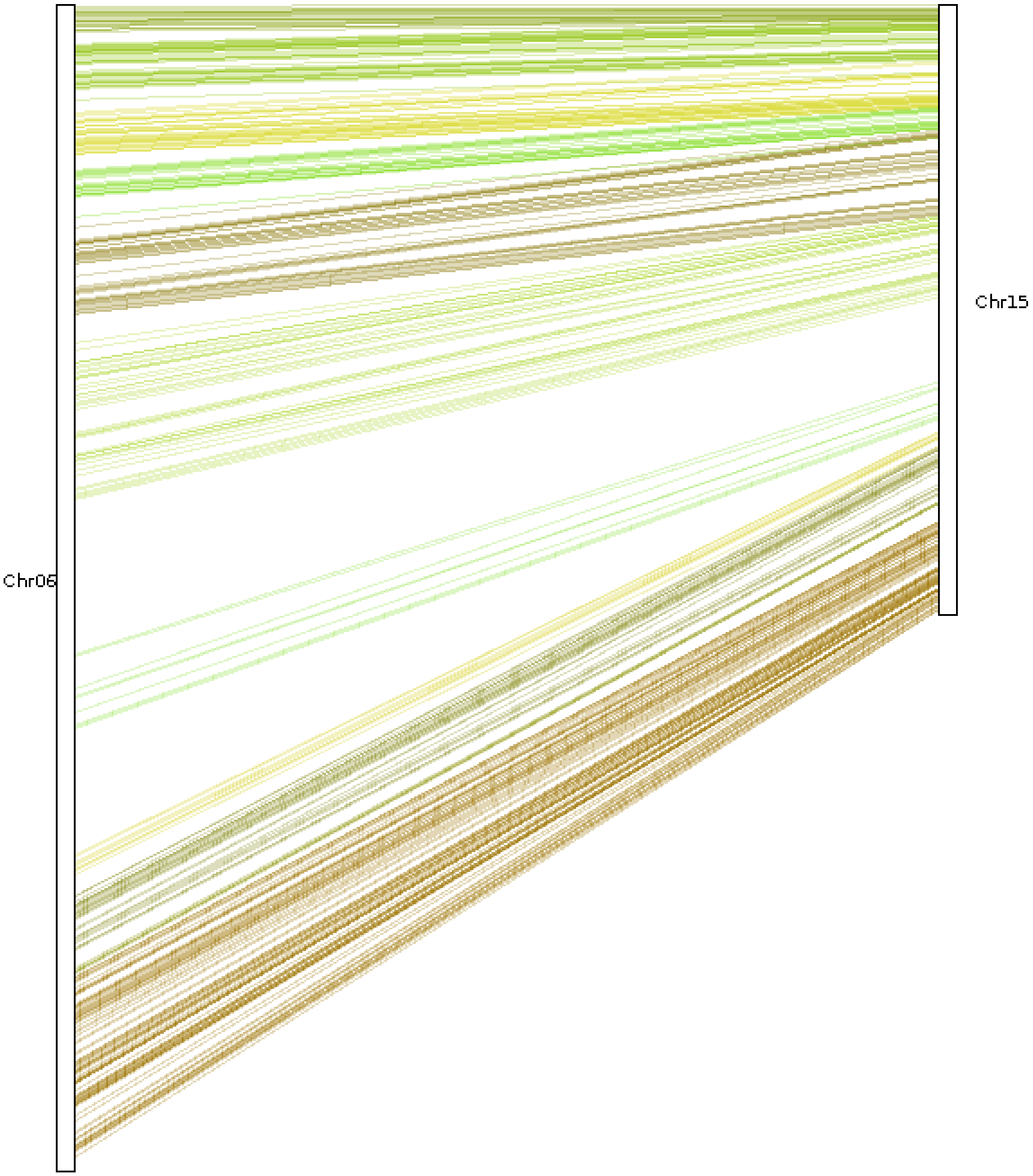

**Figure S10.** Graphical visualization of the haplotype-blocks (HB) inheritance across Chandler pedigree in the 16 chromosomes. (**A**) The inner circle highlights in grey the regions of heterozygosity and in light green the regions of homozygosity for each chromosome. The circle in the middle shows the maternally inherited HBs, while the HBs inherited from the paternal line are visualized in outer circle. In both parental line circles, missing data are highlighted in grey. Payne haplotypes are inherited along both parental lines in all chromosomes, but Chr5, Chr9, Chr10, Chr14 and Chr16. (**B**) Chandler pedigree, where Pedro is the maternal line and 56-224 the paternal line.

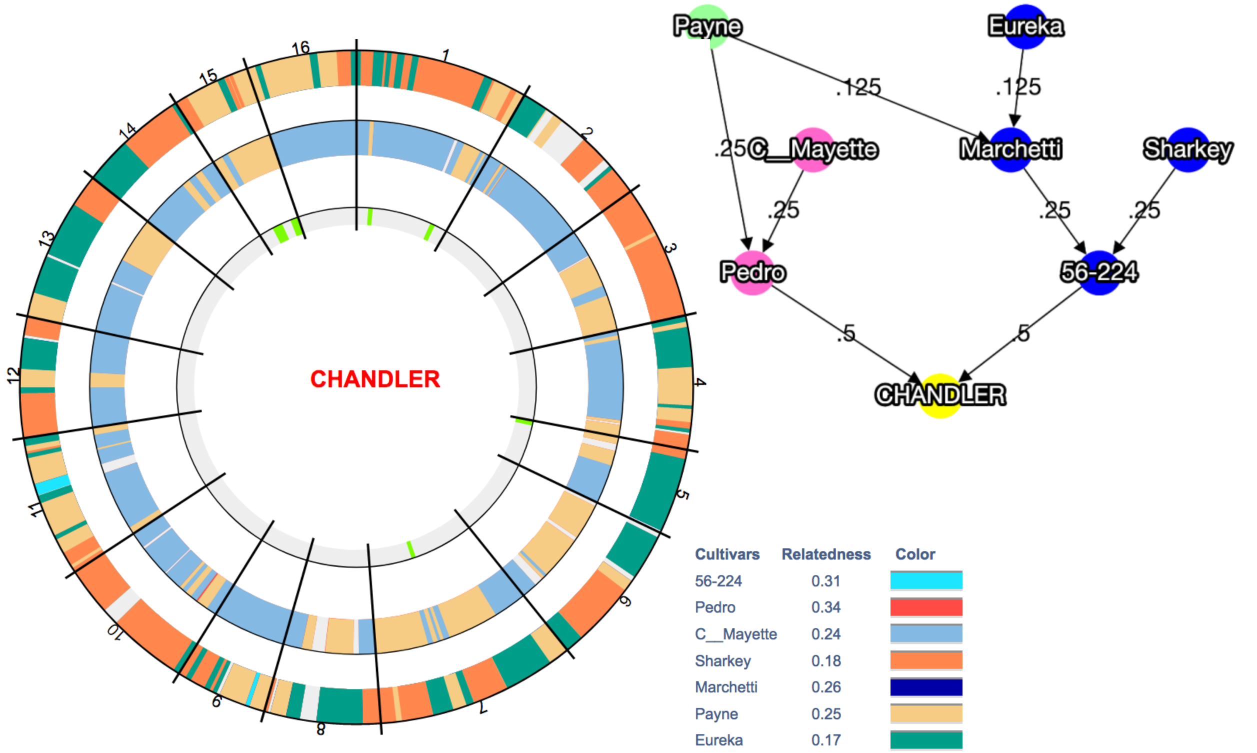

**Figure S11.** Hierarchical clustering analysis among the 23 re-sequenced founders of the UCD-WIP.

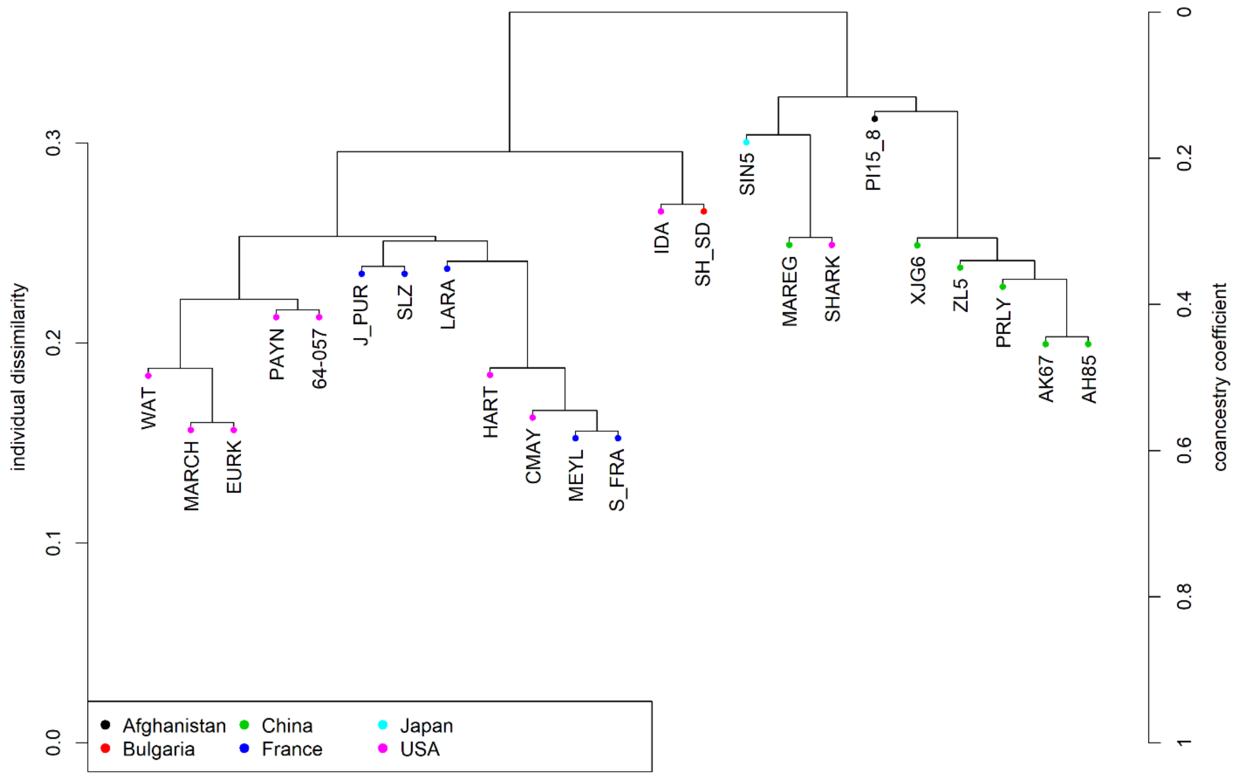

**Figure S12.** Genome scan for selective sweeps between walnuts from EU/USA and Asia. Tracks from outside to inside: (***i***) $F_{ST}$ in 500-kb windows. Windows in the 95 percentiles of the $F_{ST}$ distribution are highlighted in red; (***ii***) ROD values for 500-kb windows; (***iii***) Tajima's D in 500-kb windows for genotypes from Europe a- USA; (***iv***) Tajima's D in 500-kb windows for genotypes from Asia.

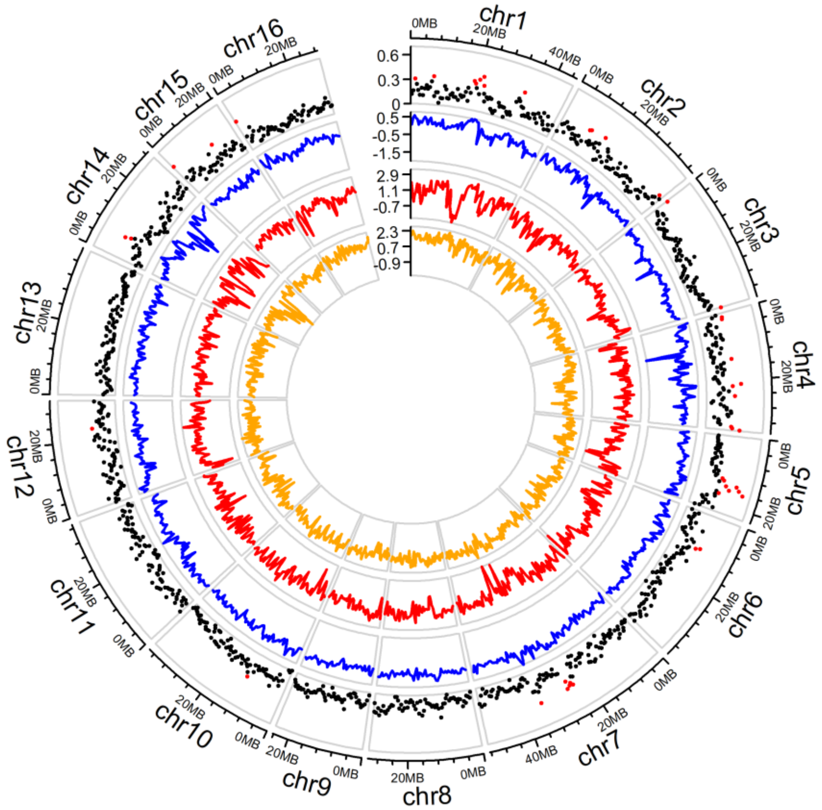

**Figure S13.** Phenotypic differences of harvesting date observed among the three genotypic classes of the marker AX-170770379 significantly associated to harvest date (Marrano et al., 2019).

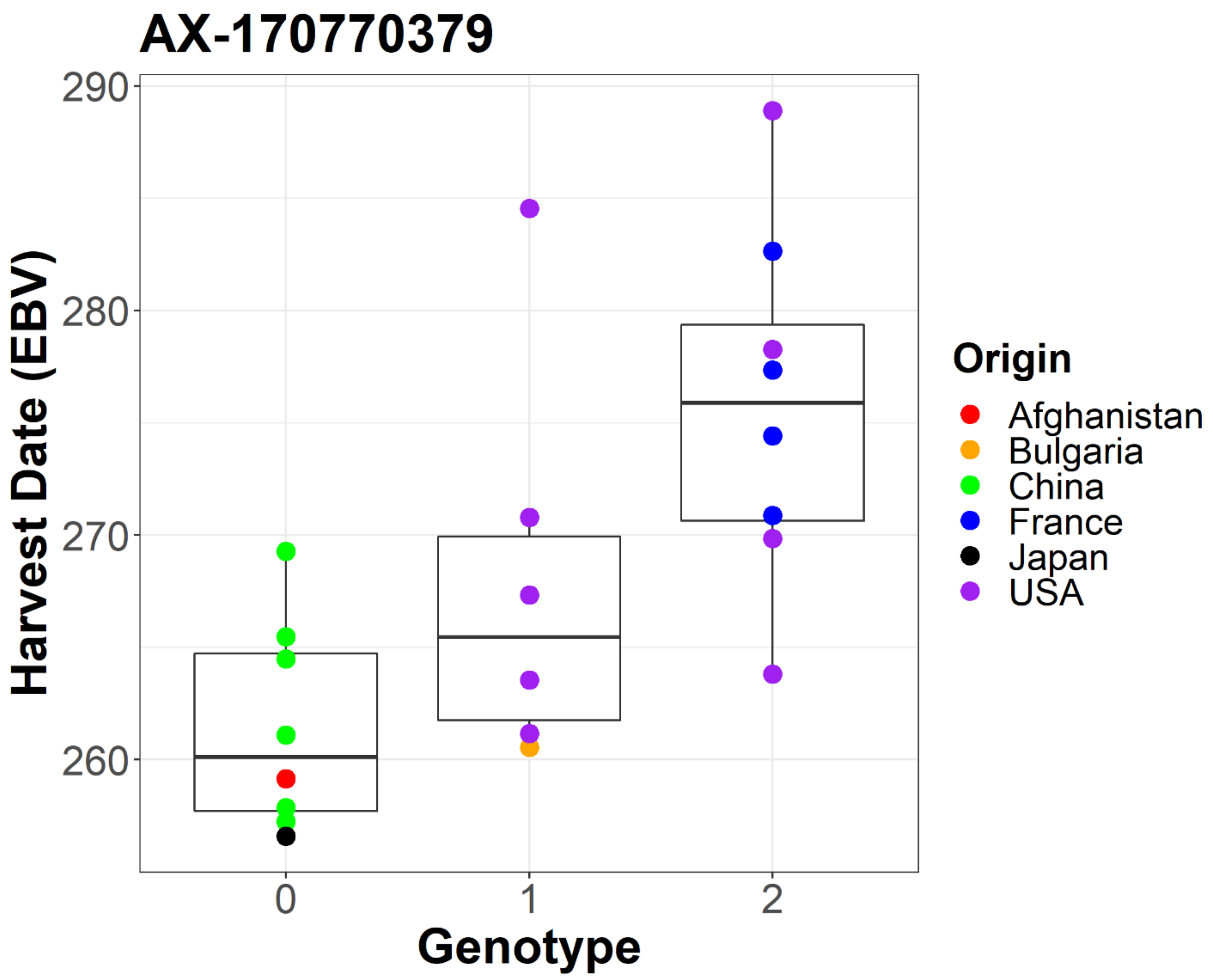
